# Epstein-Barr virus transformation creates a methionine-dependent ferroptosis vulnerability in B cells

**DOI:** 10.64898/2026.07.11.737909

**Authors:** Shaowen White, Rui Guo, Bidisha Mitra, Hanqi Li, Shunji F Li, Yifei Liao, Rishi Puri, John M Asara, Everett Stone, George Georgiou, Benjamin E Gewurz

**Affiliations:** Division of Infectious Diseases, Department of Medicine, Mass General Brigham Hospital, 181 Longwood Avenue, Boston, MA 02115, USA; Center for Integrated Solutions to Infectious Diseases, Broad Institute, Cambridge, MA 02142, USA; Harvard Medical School, Boston, MA 02115; Department of Molecular Biology and Microbiology, Tufts University, Boston, MA 02111, USA; Department of Biomedical Sciences, College of Veterinary Medicine, Cornell University, Ithaca, New York, USA; Division of Signal Transduction and Department of Medicine, Beth Israel Deaconess Medical Center, Boston, MA 02215, USA; Department of Molecular Biosciences, University of Texas, Austin, TX 78712, USA; Department of Chemical Engineering, University of Texas, Austin, TX, 78712, USA

**Author notes:** equal contribution (Shaowen White, Rui Guo).

## Abstract

Epstein-Barr virus (EBV) causes over 200,000 cancers annually, including immunoblastic lymphomas in immunosuppressed hosts. Most transformed cells arrest, yet survive when deprived of the essential amino acid methionine. We instead find that EBV transformed lymphoblastoid cell lines (LCLs), which model the EBV latency III program-driven B-cell lymphoproliferative diseases of immunosuppressed hosts, rapidly die upon methionine restriction. Methionine restriction elevated LCL lipid reactive oxygen species and triggered ferroptosis. Whereas methionine restriction hypomethylates the EBV genome and triggers viral reactivation in latency I Burkitt cells by lowering the cellular methylation potential, the LCL latency III program instead redirected methionine toward redox defense, without altering the SAM/SAH ratio. Stable-isotope tracing revealed that latency III strongly induces transsulfuration, synthesizing cysteine *de novo* to support glutathione pools. The EBV oncoprotein LMP2A, which mimics B-cell receptor signaling, supported newly infected human B cell cystathionine-β-synthase and cystathionine-γ-lyase expression and methionine dependence, phenocopied by immunoglobulin crosslinking. *In vivo*, dietary methionine restriction impaired LCL xenograft outgrowth and depleted tumor cystine. Combined methioninase and cyst(e)inase administration blocked both cysteine sources, collapsed tumor glutathione levels, and triggered ferroptosis. Our results define methionine metabolism as a targetable ferroptosis vulnerability of EBV-transformed B cells.

**Highlights:**

- Methionine restriction triggers EBV-transformed lymphoblastoid B cell ferroptosis
- EBV latency III induces transsulfuration to sustain LCL cysteine and glutathione
- Methioninase or dietary methionine restriction strongly impair LCL growth *in vivo*
- Methioninase plus cyst(e)inase collapses xenograft GSH levels and drives ferroptosis

## INTRODUCTION

Epstein-Barr virus (EBV) persistently infects over 95% of adults worldwide and is associated with over 200,000 cancers per year^1–4^. These include multiple types of lymphomas, in particular Burkitt lymphoma, Hodgkin lymphoma, and post-transplant lymphoproliferative diseases (PTLD), each of which has a distinct metabolic program that could potentially be leveraged therapeutically. EBV is also increasingly linked to the pathogenesis of human autoimmune syndromes, most notably multiple sclerosis and systemic lupus erythematosus^5–7^.

EBV pathogenesis is closely linked to the viral lifecycle, in which EBV expresses a series of latency programs to navigate the B cell compartment and that strongly remodel infected B cell metabolism states^8–12^. Newly infected primary human B cells briefly express the viral pre-latency program and then the latency IIb program, the latter of which is comprised of six Epstein-Barr nuclear antigens (EBNA) transcription factors and non-coding RNAs. These convert resting, relatively metabolically inert B cells into highly metabolically active blasts that undergo Burkitt-like hyperproliferation. The EBV oncogene EBNA2 serves as a master metabolism remodeler, together with its key target c-MYC. Over this phase of viral B-cell transformation, glycolysis, lipid metabolism and one-carbon metabolism pathways are highly induced^8–11,13,14^. One-carbon metabolism in turn supports the NADPH and glutathione biosynthesis that underpins infected B cell redox defense^8^. The methionine and one-carbon metabolism cycles are interlinked, and EBV also rapidly upregulates newly infected B cell plasma membrane methionine transporters, including SLC7A5^8^.

Beyond its role as the precursor of the universal methyl donor S-adenosyl-methionine (SAM), methionine is a source of cysteine. Following transmethylation, methionine-derived homocysteine can be condensed with serine by cystathionine-β-synthase (CBS) and cleaved by cystathionine-γ-lyase (CTH) through the transsulfuration pathway to generate cysteine *de novo*^15–19^. Cysteine is the rate-limiting substrate for glutathione (GSH) biosynthesis. Cystine is imported by the SLC7A11 antiporter, sustains the GSH pools that GPX4 uses to detoxify lipid peroxides and restrain ferroptosis^20–22^. Methionine metabolism can thereby supply the machinery of redox defense, in addition to its epigenetic roles, though how EBV-transformed B cells apportion methionine between these fates has remained undefined.

Infected B cells subsequently switch to the latency III program, which adds the EBV oncogenes LMP1 and LMP2A, mimics of CD40 and immunoglobulin B cell receptor signaling, respectively, to the EBNAs and non-coding RNAs^23–25^. This program converts infected B cells into lymphoblastoid cell lines (LCL), which model the EBV latency III–driven, immunosurveillance-dependent B-cell lymphoproliferations of immunosuppressed hosts, including PTLD and EBV+ immunoblastic diffuse large B-cell lymphoma^3,4,26–28^. Such tumors are frequently refractory to reduction of immunosuppression and to available immunotherapies, underscoring the need for approaches that exploit their altered metabolism. Although methionine metabolism is essential for EBV-driven B-cell immortalization, in part by supporting viral oncogene expression^29^, whether and how EBV-transformed cells depend on methionine for growth and survival, and the metabolic routes underlying that dependence, have remained unexplored.

In immunocompetent hosts, immune pressure limits lymphoblastoid EBV+ B-cell outgrowth, and infected cells differentiate into memory B cells, the reservoir for lifelong infection. Proliferating EBV+ memory cells express the latency I program, in which the genome-tethering protein EBNA1 is the only viral protein expressed. Most EBV+ Burkitt lymphomas likewise exhibit latency I, in which DNA methylation silences the remaining viral genes^30–32^. Consequently, methionine metabolism, which supplies SAM for cellular methylation reactions including DNA methylation, is essential for the maintenance of latency I, and dietary methionine restriction or DNA hypomethylation de-represses a range of EBV genes within Burkitt tumor xenografts, although effects on Burkitt growth and survival were not defined^29^. Whether latency III cells share this epigenetically-rooted methionine dependence, or instead rely on methionine for distinct roles, was unknown.

Metabolic remodeling is a hallmark of cancer, motivating interest in targeting altered tumor metabolism. EBV dynamically sensitizes newly infected cells to ferroptosis, a form of regulated cell death in which iron-catalyzed lipid reactive oxygen species (ROS) drive catastrophic phospholipid membrane damage^21,33–38^. Whereas newly infected cells undergoing Burkitt-like proliferation, and Burkitt tumor cells themselves, are highly sensitive to ferroptosis induced by cystine-import blockade, lymphoblastoid cells become progressively resistant as they evolve into LCLs^37^. The pathways underlying this acquired LCL ferroptosis resistance remain poorly understood, particularly those interconnecting with methionine metabolism, which has recently been implicated in ferroptosis sensitivity^39^. This raises the possibility that methionine restriction could reverse LCL redox protection.

Nearly all cancers undergo growth arrest upon withdrawal of the essential amino acid methionine, even when it is replaced by its metabolic precursor homocysteine. This methionine addiction is known as the Hoffman effect (also termed the methionine dependence, or methionine stress sensitivity, of cancer)^40–42^. By contrast, non-transformed cell proliferation is restored by homocysteine supplementation under methionine restriction. The Hoffman effect is thought to reflect an increased cancer-cell demand for methionine-derived metabolites^40^.

Given the bidirectional links between EBV latency and methionine metabolism, we asked whether LCLs undergo the Hoffman effect upon methionine withdrawal. Intriguingly, methionine restriction instead rapidly triggered LCL death *in vitro*. We identify EBV oncogene–driven roles for methionine metabolism in cysteine synthesis via transsulfuration, in cystine metabolism, and in support of NADPH and glutathione pools that together restrain ferroptosis. Dietary methionine restriction or methioninase administration significantly impaired LCL xenograft outgrowth *in vivo*, and combined methioninase and cyst(e)inase administration strongly depleted xenograft glutathione and triggered ferroptosis.

## RESULTS

### The EBV latency III program dictates methionine metabolism dependency for survival

To gain insights into how EBV latency programs affect B cell dependency on extracellular methionine, we cross-compared methionine restriction effects on two tumor-derived EBV+ Burkitt B cells versus LCLs. Burkitt cells typically have the EBV latency I program, whereas LCLs express EBV latency III (**Figure 1A**). Methionine restriction triggered relatively low levels of cell death, marked by plasma membrane Annexin V and 7-AAD uptake, in EBV+ Burkitt P3HR-1 or MUTU I cells over the first 48 hours of methionine restriction (defined as growth in 1μM methionine versus the 100μM methionine typically present in RPMI). By contrast, 40-60% of GM12878 or GM13111 LCLs underwent cell death when grown under methionine restriction at this early timepoint (**Figure 1B, S1A-B**). By 72 hours of methionine restriction, >80% of LCLs lost viability, as compared with a lower percentage of EBV+ Burkitt cells (**Figure 1B**). Methionine restriction increased G2-phase cell cycle arrest in surviving LCLs (**Figure S1C-D**).

**Figure 1.**
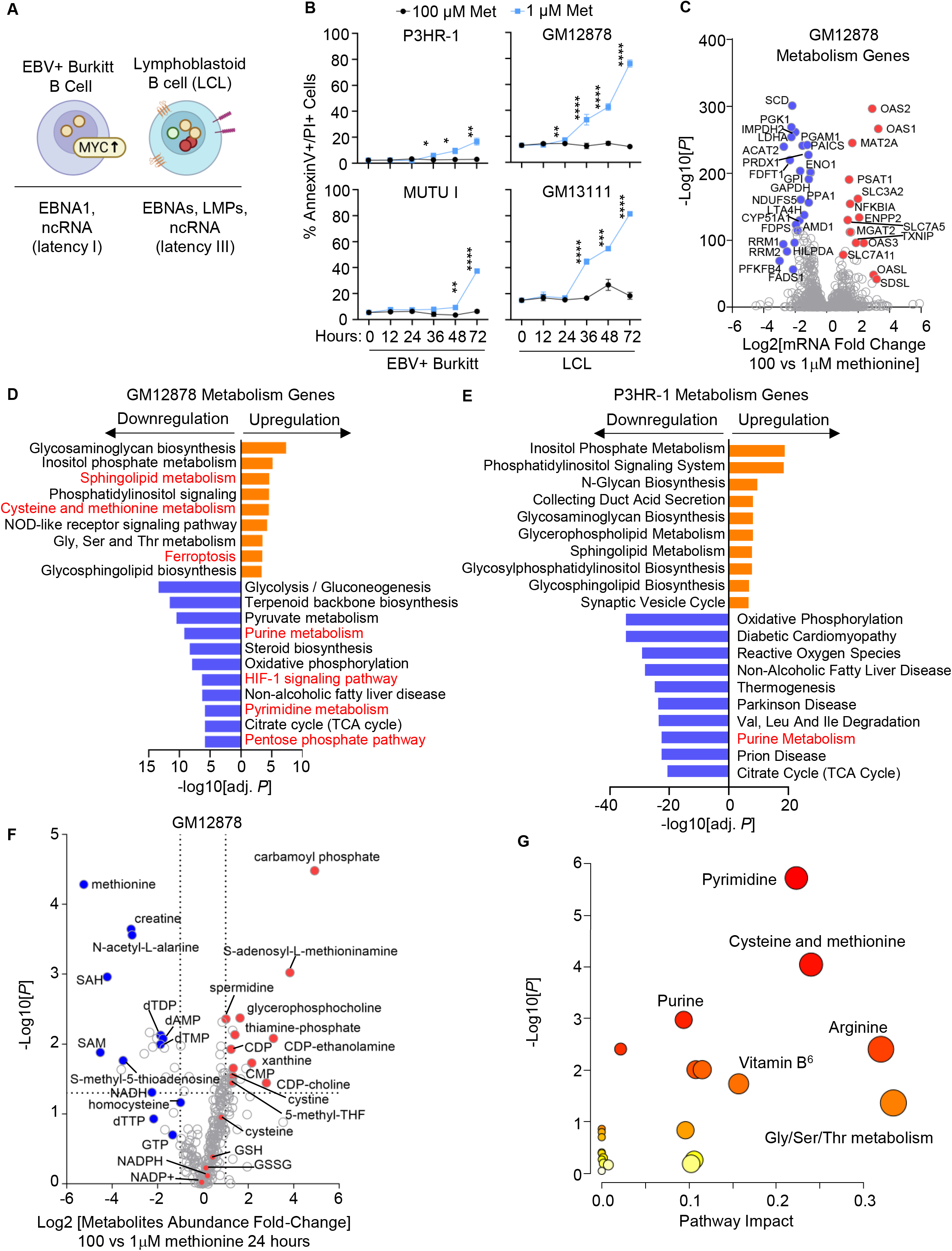
Differential methionine restriction effects on EBV-transformed LCLs vs Burkitt cells. (A) EBV+ Burkitt cells typically use the latency I program and have elevated c-MYC, while LCL use the latency III program. (B) Methionine restriction effects on Burkitt vs LCL survival. Shown are mean ± SD percentage of 7-AAD+/Annexin V+ (dead) cells from n=3 replicates. EBV+ Burkitt (P3HR-1, MUTU I) and LCLs (GM12878, GM13111) were seeded in RPMI 1640 with 100 or 1μM methionine (Met) for the indicated times. Student’s T-test, ****p<0.0001, ***p<0.001, **p<0.01, *p<0.05. (C) Methionine restriction effects on LCL metabolism gene^89^ expression. Volcano plot from n=3 RNAseq replicates of GM12878 grown in media with 100 vs 1μM methionine for 30 hours. (D) Top 10 Enrichr KEGG pathways enriched within differentially expressed genes in LCLs grown in 100 vs 1 μM methionine, as in (C), using an adjusted P-value <0.05 cutoff. (E) Top 10 Enrichr KEGG pathways enriched within differentially expressed genes in P3HR-1 Burkitt B cells grown in 100 vs 10 μM methionine for 72 hours, using an adjusted P-value <0.05 cutoff. Data are from^29^. (F) Methionine restriction effects on the LCL metabolome. Volcano plot from LC-MS polar metabolite analysis from n=3 LC-MS replicates of GM12878 grown in media with 100 vs 1μM methionine for 30 hours. (G) MetaboAnalyst analysis of methionine restriction driven metabolic pathway impact from the data presented in (F). Higher pathway impact values indicates stronger effects of methionine restriction on the indicated pathway.

In order to identify key methionine restriction effects on GM12878 LCLs, which are a Tier 1 Encode project cell line^43^, we next performed RNAseq analysis at 24 hours of growth in methionine restricted versus replete media, a timepoint prior to the onset of high levels of cell death. Methionine restriction significantly altered metabolism gene expression, with pentose phosphate pathway (PPP) and nucleotide metabolism amongst the most highly downmodulated metabolic pathways (**Figure 1C-D, Table S1**). For instance, PFKFB4, which increases glycolysis flux towards PPP, was amongst the most highly downmodulated LCL gene, and multiple glycolysis pathway genes were similarly downmodulated, including ENO1 and GAPDH. TXNIP, which negatively regulates glucose uptake^44^, was instead upregulated. Similarly, the nucleotide metabolism gene IMPDH2, which we recently found to be essential for LCL survival by supporting *de novo* purine biosynthesis downstream of EBV oncogene LMP1^45^, was highly downmodulated by methionine restriction (**Figure 1C, Table S1**).

Methionine restriction highly upregulated multiple LCL metabolic pathways, including cysteine and methionine metabolism, as well as ferroptosis (**Figure 1C-D**). Interestingly, multiple interferon-stimulated genes, including OAS1-3 and OASL were upregulated by methionine restriction, suggesting unexplored cross-talk between LCL methionine metabolism and innate immune signaling. However, methionine restriction did not significantly change abundances of EBV transcripts other than EBNA3C and EBER2, which were reduced by approximately 4-fold at this early timepoint (**Figure S2A**). Cross-comparison of these results with published transcriptomic analyses ^29^ highlighted that methionine restriction altered a partially overlapping set of EBV+ Burkitt B cell metabolism genes (**Figure 1E, S2B-F**). Methionine restriction highly downmodulated Burkitt oxidative phosphorylation and reactive oxygen species pathways and upregulated multiple lipid metabolism pathways, only a subset of which were upregulated in LCLs by methionine restriction (**Figure 1E, S2E**).

We next performed liquid chromatography-mass spectrometry (LC-MS) polar metabolite analysis to define methionine restriction effects on the LCL metabolome at the 24-hour timepoint. As expected, methionine was the most depleted metabolite from LCLs grown in 1μM methionine, with levels depleted by ∼30-fold (**Figure 1F, Table S2**). This result highlights the extent to which LCLs import and rapidly metabolize methionine. Similarly, the methionine cycle metabolites S-adenosyl-methionine (SAM) and S-adenosyl-homocysteine (SAH) were rapidly depleted, as were multiple nucleotides, likely reflecting interconnecting LCL methionine cycle and one-carbon metabolism (**Figure 1F-G**). However, methionine restriction instead increased LCL cytidine monophosphate (CMP) and diphosphate (CDP), possibly as a result of altered salvage metabolism. However, the SAM/SAH ratio, which defines the cellular methylation potential, was not significantly changed. This result contrasts with methionine restriction effects on EBV+ Burkitt cells, which strongly reduces the SAM/SAH ratio, resulting in DNA hypomethylation and induction of EBV latency and lytic gene expression^29^. Taken together, these results highlight distinct transcriptomic and metabolomic responses to methionine restriction in EBV-transformed LCL versus Burkitt B cells.

### Methionine restriction elevates lymphoblastoid B cell lipid ROS levels

We next characterized the LCL death pathway triggered by methionine restriction. LCL culture in media with 1μM methionine modestly increased executioner caspase 3/7 activity and stimulated a low level of PARP1 cleavage (**Figure 2A, S3A**). However, treatment with the pan-caspase inhibitor zVAD-FMK did not rescue methionine restriction-driven LCL death, suggesting that it did not trigger apoptosis (**Figure S3B**). Likewise, necroptosis pathway inhibition by the RIPK1 inhibitor Nec-1^46^ did not rescue methionine restriction-induced LCL death (**Figure S3B**). We next measured methionine restriction effects on Burkitt versus LCL lipid reactive oxygen species (ROS) levels, which are elevated in cells triggered for ferroptosis. Methionine restriction significantly increased LCL, but not Burkitt lipid ROS levels, as judged by BODIPY C11 staining (**Figure 2B**). Interestingly, homocysteine supplementation rescued methionine restriction induced LCL death, further differentiating LCL methionine dependence from the Hoffman effect (**Figure 2C**).

**Figure 2.**
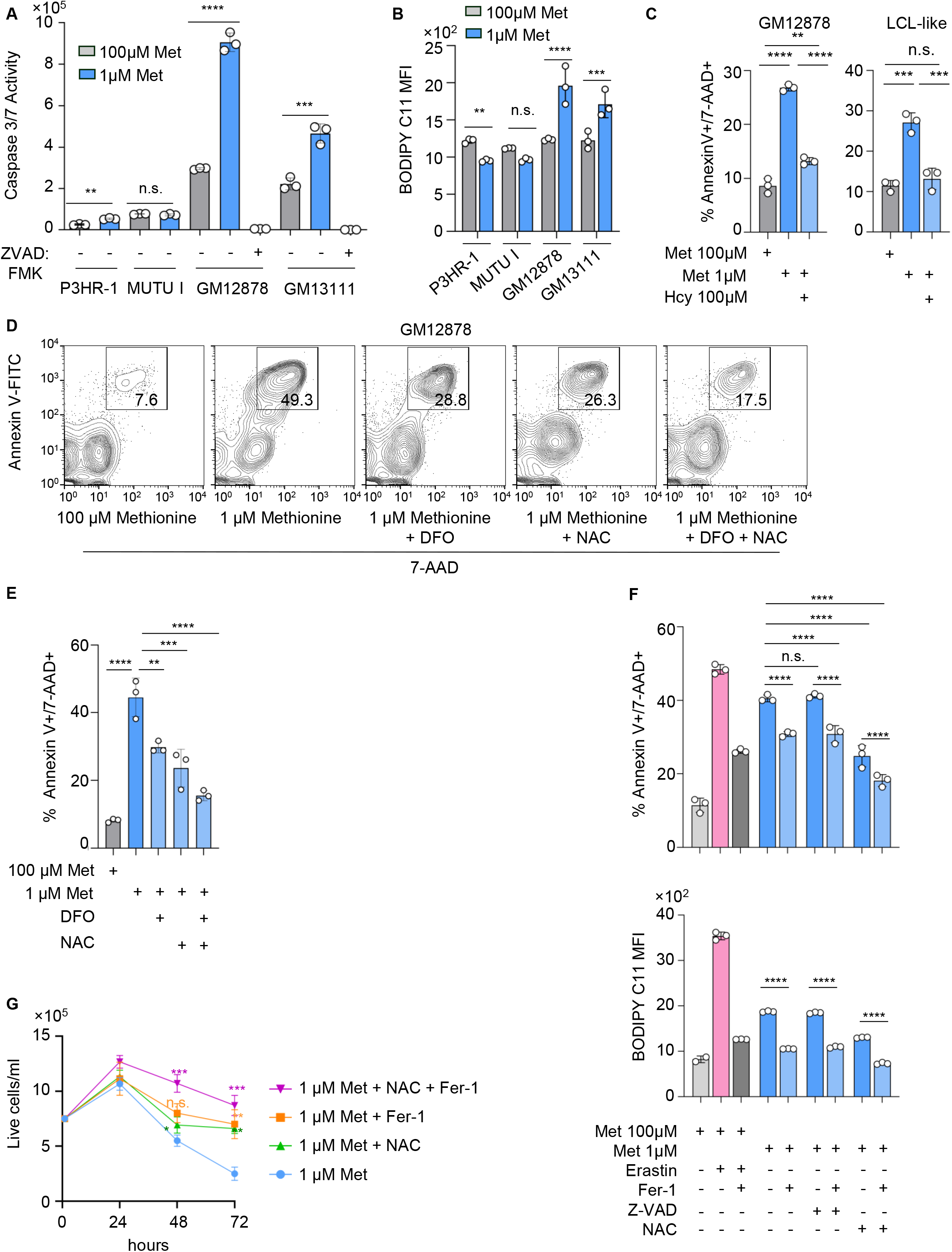
Methionine restriction triggers LCL ferroptosis. (A) Methionine restriction effects on LCL caspase 3/7 activity. Mean ± SD caspase 3/7 activity from n=3 replicates of the indicated cells cultured in media with 100 versus 1μM methionine, without or with caspase-inhibitor Z-VAD-FMK (ZVAD, 20μM) for 30 hours. (B) Methionine restriction effects on LCL lipid ROS. Mean ± SD BODIPY C11 levels in cells cultured as in (A). (C) Homocysteine supplementation effects on cell death caused by methionine restriction in LCL. Shown are mean ± SD percentage of 7-AAD+/Annexin V+ (dead) cells from n=3 replicates. GM12878 or LCL-like cells at approximately 3 months post-EBV transformation were grown in RPMI 1640 with 100 or 1μM Met, with 100 μM DL-homocysteine thiolactone hydrochloride (Hcy) supplementation as indicated for 30 hours. (D) Iron chelation and antioxidant effects on methionine-restriction driven LCL death. FACS analysis of annexin V positivity and 7-AAD uptake in GM12878 LCLs cultured in media with 100 vs 1μM methionine and treated with deferoxamine (DFO, 100μM) and/or N-acetyl cysteine (NAC, 10mM) for 48 hours. Plots are representative of n=3 replicates. (E) Mean ± SD dead cell percentage (7-AAD+/Annexin V+) from n=3 replicates of cells treated as in (D). (F) Methionine restriction causes LCL ferroptosis. Mean ± cell death percentage (top) versus BODIPY C11 (lipid ROS) mean fluorescence intensity (MFI, bottom) levels from n=3 replicates of GM12878 cultured in media with 100 vs 1μM methionine and treated with ferrostatin 1 (Fer-1, 5μM), NAC (10mM) and/or Z-VAD (20μM) for 30 hours, as indicated. As a + control, cells were treated with erastin 1μM for 18 hours. (G) Antioxidant rescue of methionine restriction driven LCL death. Mean ± SD live cell numbers from n=3 replicates of GM12878 cultured in media with 1μM methionine and treated with Fer-1 and/or NAC as indicated. Student’s t-test was used for (A). One-way ANOVA was used for (C, E, G). Two-way ANOVA was used for (F). For all statistical analyses, ****p<0.0001, ***p<0.001, **p<0.01, *p<0.05.

To further test the hypothesis that methionine restriction triggers LCL ferroptosis, we treated GM12878 LCLs grown under methionine restriction for 48 hours with the iron chelator deferoxamine (DFO) and/or with N-acetyl-cysteine (NAC), a cysteine prodrug that supports glutathione biosynthesis and that scavenges ROS to inhibit ferroptosis. DFO or NAC treatment significantly rescued cell viability, as measured by FACS analysis of 7-AAD/Annexin V stained cells. Combined DFO/NAC treatment nearly completely blocked methionine restriction-induced LCL death (**Figure 2D-E**). Similar results were found with GM13111 LCLs (**Figure S3C-D**). In further support of LCL ferroptosis induction, the lipid ROS detoxifying antioxidant Fer-1 also reduced methionine restriction driven lipid ROS and LCL death, to a somewhat greater extent when combined with NAC (**Figure 2F-G**).

Although DFO and NAC treatment also decreased methionine restriction triggered caspase activity, combined zVAD-FMK + Fer-1 treatment did not reduce LCL death more than treatment with Fer-1 alone (**Figure 2F, S3E**). Similar results were obtained in GM13111 LCLs (**Figure S3F-G**). However, methionine restriction did not increase overall intracellular oxidant levels in either GM12878 or GM1311 LCLs, as judged by FACS analysis of H2DCFCDA stained cells (**Figure S3I**). Likewise, the ROS scavengers Trolox (50μM) or TEMPO (125μM) ^47,48^ did not rescue LCLs from methionine restriction driven cell death, even though they successfully blocked H_2_O_2_-triggered ROS production (**Figure S3J-K**). Taken together, these results suggest that methionine restriction predominantly kills LCLs through ferroptosis induction, which is highly unusual for cancer cells.

### Methionine metabolism supports newly EBV infected B cell redox defense

EBV induces lipid metabolism^8^ and dynamically sensitizes newly infected primary human B cells to ferroptosis, particularly at early timepoints post-infection^37^. We therefore next tested the hypothesis that newly EBV-infected cells are dependent on methionine metabolism for survival at key stages of B-cell transformation into LCLs. To do so, we isolated human peripheral blood primary B cells by negative selection and infected them with the EBV B95.8 strain at a multiplicity of infection of 0.3, an optimal dose for EBV-driven B-cell outgrowth^49^. Newly infected cells were cultured in media with 100μM methionine, and parallel cultures were shifted to media with 10μM methionine, starting at days 0, 2, 5 or 10 days post-infection for two or four days (**Figure 3A**). We used 10uM methionine as the methionine restriction dose for primary peripheral blood B cells based on our prior study, which found that this dose strongly impaired EBV-mediated B-cell transformation^29^.

**Figure 3.**
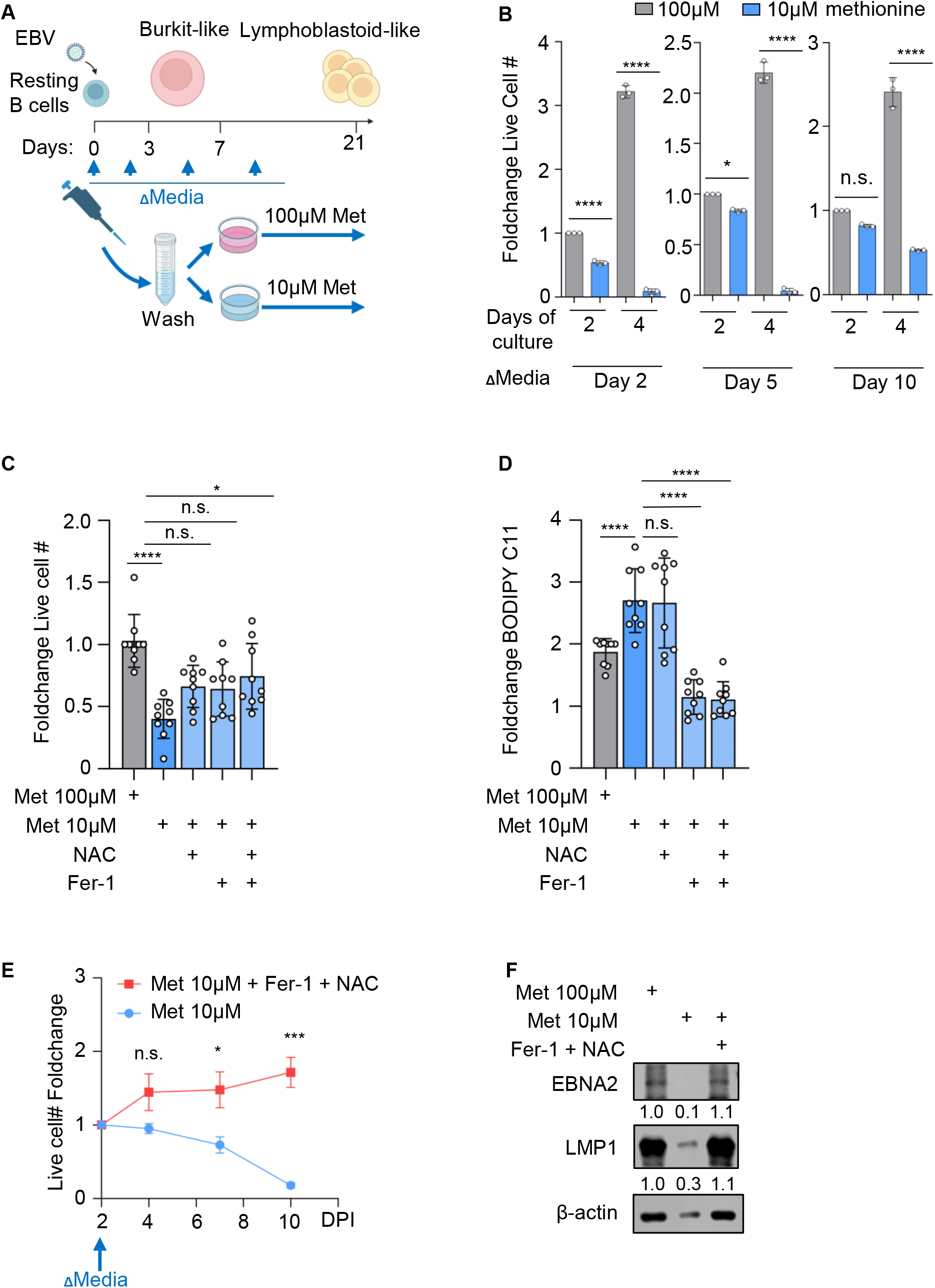
EBV induces methionine dependence in newly-infected human peripheral blood B cells. (A) Experimental design for test of methionine dependence at key timepoints of EBV-mediated primary B cell transformation. (B) Methionine restriction effects on primary B cell outgrowth. B cells were infected by EBV strain B95.8 and then cultured in media with 100μM Met for 2, 5, or 10 days. Cultures were then split into media with 100 vs 10μM Met for 2 or 4 days. Shown are mean ± SD fold-change of live B cell numbers, normalized by cell counts in 100μM Met at two days post-media change. (C) Methionine restriction antioxidant rescue effects on newly infected B cells. Mean ± SD foldchange live cell numbers from n=3 donors (3 replicates each). Naïve B cells were cultured in media with 100uM Met until 2 days post infection and then cultured in media with 100 vs 10μM Met. Fer-1 and/or NAC were added at day 2 post-infection, as indicated. (D) Antioxidant effects on methionine restriction induced lipid ROS in newly infected B cells. Mean ± SD fold-change BODIPY MFI of cells cultured as in (C). (E) Antioxidant effects on newly infected, methionine restricted B cells. Mean ± SD foldchange live cell numbers of primary B cells from n=3 donors infected by EBV and cultured in media with 100μM Met for two days, then shifted into media with 10μM met ± Fer-1 and NAC. (F) Antioxidant rescue of EBV oncogene expression in newly infected methionine restricted B cells. Representative immunoblots of WCL from cells cultured for seven days in media with 100μM Met, versus in 10μM Met-/+ Fer-1 and NAC. Student’s t-test was used in B and E. One-way ANOVA was used in (C-D). ****p<0.0001, ***p<0.001, **p<0.01, *p<0.05.

Methionine restriction strongly reduced EBV-transformed B cell outgrowth when begun at Day 2 post-infection, a timepoint prior to the first infected cell mitosis where EBV significantly increases cell volume and drives B-cell remodeling in preparation for rapid B-cell outgrowth. Similarly, methionine restriction significantly impaired outgrowth of cells when begun at Day 5 post-infection, in the middle of the period of EBV-driven Burkitt-like B-cell hyperproliferation (**Figure 3A-B**). We observed significant but somewhat less impairment of EBV-driven B cell outgrowth when methionine restriction was initiated at Day 10 post-infection, an early timepoint of the infected cell lymphoblastoid phase (**Figure 3A-B**).

To then investigate the extent to which newly EBV-infected cells are dependent on abundant extracellular methionine for survival, we cross-compared the percentage of live B cells cultured in 100 vs 10μM methionine, starting at Day 2 post-infection and continued for 2 days. Methionine restriction significantly decreased live cell number, as measured by vital die uptake analysis, and this could be partially rescued by addition of either NAC or Fer-1 (**Figure 3C**). Consistent with ferroptosis induction, methionine restriction increased BODIPY C11 levels, and this methionine restriction-induced increase could be blocked by addition of Fer-1, though interestingly not by NAC (**Figure 3D**). These results are consistent with a model in which methionine restriction drives lipid-ROS production and ferroptosis induction in newly infected cells, and also induces ROS-driven caspase activity in parallel.

We next tested the extent to which Fer-1 and NAC addition could rescue newly infected B-cell outgrowth. Peripheral blood B cells were infected and grown for two days in media with 100μM methionine. At Day 2 post-infection, cells were washed and plated in media with 10μM methionine, in the absence or presence of Fer-1 + NAC. While cells grown in the absence of Fer-1 + NAC rapidly died, Fer-1 + NAC treated cell number was relatively constant over 8 days of culture (**Figure 3E**). Thus, blocking ferroptosis converted methionine-restriction-driven death into growth arrest in newly infected B cells.

We previously reported that methionine restriction diminishes EBV oncogene expression at this early timepoint of 2 days post-infection^49^. To further examine the mechanism by which Fer-1+NAC rescues cell survival, we measured their effect on the expression of EBNA2 and LMP1, key EBV oncogenes necessary for B cell immortalization. As previously reported, methionine restriction strongly impaired both EBNA2 and LMP1 expression at this timepoint. Interestingly however, Fer-1+NAC rescued EBNA2 and LMP1 expression (**Figure 3F**), suggesting that lipid ROS may block EBV oncogene expression in newly EBV-infected primary human B cells.

### EBV oncogene-driven methionine metabolism supports intracellular cysteine pools

Cysteine plays key roles in redox defense, including as the rate-limiting amino acid of glutathione biosynthesis^50^. Furthermore, LCLs are dependent on GPX4 and glutathione mediated lipid ROS detoxification to prevent ferroptosis induction^37^. We therefore hypothesized that a key LCL methionine metabolism role is to support intracellular cysteine pools, including by the transsulfuration pathway, which synthesizes cysteine *de novo* from methionine and serine^15–18^. To test this hypothesis, we performed GM12878 LCL stable isotope tracing using [U-¹³C_5_] methionine, where each methionine carbon was ^13^C-labeled (hereafter referred to as m+5). GM12878 LCLs were labeled for six hours, followed by LC-MS analysis. In support of robust LCL methionine uptake and metabolism, >50% of the SAH pool was isotopically labelled, most of which with four (m+4) carbon labels (**Figure 4A, Table S3**). The methionine cycle metabolizes SAH to homocysteine, which can be used to regenerate methionine or which together with serine can be condensed by the transsulfuration pathway enzyme CBS to make cystathionine (**Figure 4A**). We observed robust cystathionine ¹³C labeling, and ¹³C-labeled SAH and cystathionine ion intensities were similar (**Figure 4A, Table S3**). Together, our tracing suggests that CBS, which is both transcriptionally and post-translationally regulated, is highly active in LCLs in support of transsulfuration.

**Figure 4.**
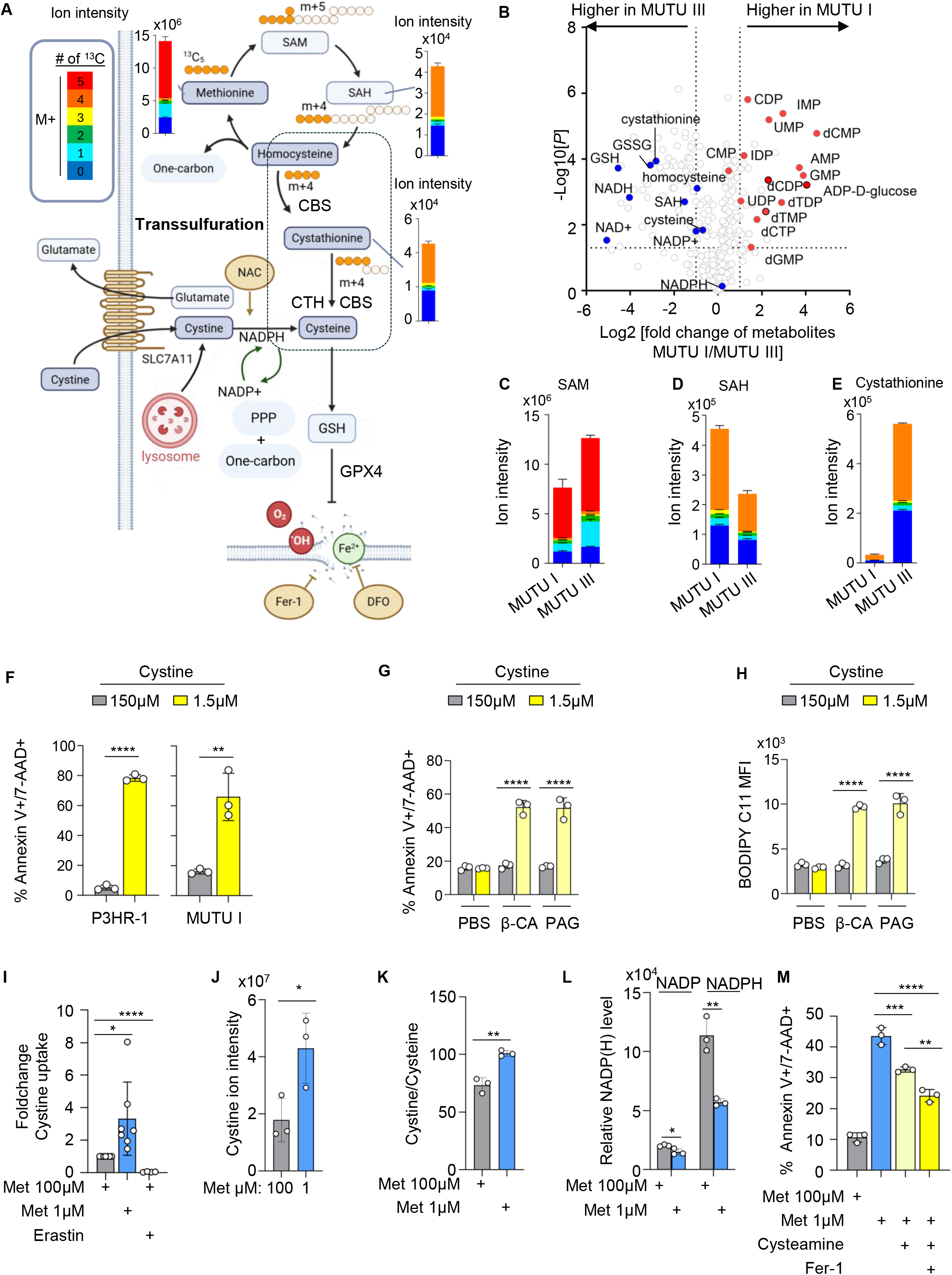
EBV Latency III induces transsulfuration to support lymphoblastoid B cell cysteine pools. (A) U^13^C-methionine metabolic tracing in LCLs. Schematic showing metabolic tracing of U^13^C-methionine into cystathionine by the transsulfuration pathway. Orange circles denote heavy ^13^C atoms, while white circles denote light ^12^C atoms. Overlaid on the schematic are SAH and cystathionine ion intensities in metabolite extracts from 10 million GM12878 LCLs fed U^13^C-methionine for six hours prior to LC-MS analysis. Data show the mean from n=3 replicates. (B) EBV latency III program metabolome remodeling. Volcano plot from LC-MS polar metabolite analysis from n=3 LC-MS replicates of isogenic MUTU I versus MUTU III Burkitt cells, which differ by EBV latency I versus III programs, respectively. (C-E) EBV latency III effects on methionine metabolites. Mean SAM (C), SAH (D) and cystathionine (E) ion intensities in metabolite extracts of 10 million MUTU I vs III cells fed U^13^C-methionine for six hours prior to LC-MS analysis of n=3 replicates. Cells were grown in media with 100μM methionine. Isotopologue color scheme as in (A). (F) Cystine restriction effects on EBV+ Burkitt viability. Mean ± SD 7-AAD+/Annexin V+ percentages from n=3 replicates of P3HR-1 and MUTU I cells cultured in media with 150 versus 1.5μM cystine for 36 hours. (G) Transsulfuration inhibition effects on LCL sensitivity to cystine restriction. Mean ± SD 7-AAD+/Annexin V+ percentages from n=3 replicates of GM12878 LCLs cultured in media with 150μM or 1.5μM cystine, in the presence of PBS vehicle, the CBS inhibitor β-cyano-L-alanine (βCA, 100μM) or the cystathionine-γ-lyase inhibitor DL-propargyl glycine (PAG, 1mM) for 36 hours. (H) Transsulfuration inhibition effects on cystine restriction induced LCL lipid ROS. Mean ± SD BODIPY C11 levels in cells treated as in (G). (I) Methionine restriction effects on LCL cystine uptake. Mean ± SD foldchange of cystine uptake in levels from n=3 replicates of GM12878 cultured in media with 100 or 1μM Met for 30 hours, or as a control grown in 100μM Met and treated with erastin (1μM) for 18 hours. Cystine uptake was determined by a platereader luminescence assay. (J) Methionine restriction effects on LCL cystine level. Mean ± SD relative cystine levels from the GM12878 shown in (F). (K) Methionine restriction effects on LCL cystine/cysteine ratio. Mean ± SD relative cystine/cysteine ratio from the GM12878 shown in (F). (L) Methionine restriction effects on LCL NADP(H). Mean ± SD relative NADP(H) levels in GM12878 cultured in media with 100 or 1μM Met, as measured by a NADP(H) luminescence assay. (M) Cysteamine effects on methionine restriction induced LCL death. Mean ± SD 7-AAD+/ Annexin V+ percentages from n=3 replicates of GM12878 cultured in media with 100 vs 1μM Met, with Fer-1 and/or the cystine-reducing aminothiol cysteamine (100μM), as indicated. Student’s t-test was used for statistical analysis in F, J, K, L. One-way ANOVA was used for (G, H, I, M). ****p<0.0001, ***p<0.001, **p<0.01, *p<0.05.

We next hypothesized that EBV latency programs modulate methionine metabolism to support cysteine abundance and downstream glutathione pools. To gain further insights, we therefore first performed LC/MS analysis on isogenic EBV+ Burkitt lymphoma B cells, which differ only by EBV latency I vs III programs (MUTU I vs III)^51^. In support of this hypothesis, the methionine metabolites SAH and cystathionine were higher in MUTU III cells, with cystathionine nearly 8-fold more abundant in MUTU III (**Figure 4B, Table S4**).

To then more directly measure methionine flux, we cross-compared [U-¹³C₅] methionine isotope tracing of MUTU I versus III cells. Cells were again [U-¹³C_5_] methionine labeled for 6 hours and then profiled by LC-MS. A roughly similar percentage of MUTU I and III SAM pools were isotopically labeled, with most having M+4 carbon labels (**Figure 4C, Table S3**). Though SAH levels were lower in MUTU III, a similar percentage of the SAH pool was isotopically labeled in both MUTU I and III cells (**Figure 4D, Table S3**). However, consistent with robust latency III program transsulfuration induction, LC-MS analysis revealed a much larger cystathionine pool in MUTU III than I, approximately half of which was M+4 labelled (**Figure 4E, Table S3**). Taken together, these data suggest that the EBV latency III program induces CBS activity and the transsulfuration pathway.

Cysteine levels were approximately 2-fold higher in MUTU III than I (**Figure 4E**). We therefore next tested cystine restriction effects on Burkitt cells versus LCLs. Interestingly, whereas Burkitt cells rapidly underwent cell death upon culture in media with 1.5μM cystine, LCLs tolerated cystine withdrawal for 36 hours (**Figure 4F-G**). However, the CBS inhibitor β-cyano-L-alanine (βCA) or the CTH inhibitor DL-propargylglycine (PAG) strongly sensitized LCLs to cystine restriction (**Figure 4G**). Consistent with ferroptosis induction as the cell death mechanism, cystine restriction highly induced lipid ROS levels in βCA or PAG treated LCLs (**Figure 4H**). Taken together, these data suggest that latency III-driven transsulfuration enables EBV-transformed cells to buffer against changes in extracellular cystine.

We therefore next directly measured methionine restriction effects on LCL cystine and cysteine levels. Of note, LCLs highly express the cystine/glutamate antiporter SLC7A11, which is upregulated by LMP1^22^. To define methionine restriction effects on cystine uptake, we cultured cells in media with 100 vs 1μM methionine for 30 hours. Even at this early timepoint, methionine restriction significantly increased cystine uptake, both in GM12878 and GM13111 LCLs, which could be blocked by erastin (**Figure 4I, S4A**). However, methionine restriction paradoxically increased LCL cystine levels and the cystine/cysteine ratio (**Figure 4J-K**). This result may have been produced by two methionine restriction effects: increased cystine uptake, as methionine restriction increased SLC7A11 levels by 3.3 fold (**Table S1**), and impaired transsulfuration. Furthermore, we found methionine restriction reduced LCL NADPH levels using an orthogonal assay (**Figure 4L, S4B**), likely due to impairment of the pentose phosphate and one-carbon metabolism pathways (**Figure 1D**). Treatment with the cystine-reducing aminothiol cysteamine diminished methionine restriction driven LCL death (**Figure 4M**). Taken together, these results support a model in which methionine metabolism supports LCL intracellular cysteine pools through transsulfuration, by increasing cystine uptake and by supporting its NADPH-dependent reduction to cysteine.

### EBV oncogene LMP2A as a driver of transsulfuration and methionine dependence

To gain insights into specific EBV oncogene effects on infected B cell methionine survival dependence, we utilized a panel of EBV wildtype versus oncogene knockout (KO) viruses. Across the panel, one of each of the eight latency III oncogenes is KO^49^. Each virus encodes a GFP marker, enabling precise normalization of infectious EBV titers by the green Raji assay^52^. Human peripheral blood B cells from n=3 donors were isolated by negative selection and infected with wildtype B95.8 EBV, or with an EBV latency III oncogene knockout. Cells were grown for two days in methionine replete media, and then split into parallel cultures grown in methionine replete versus restricted media for three additional days. Relative live cell numbers were calculated at the end of the assay. We note that most EBNA2 KO infected B cells were not viable as previously published^53,54^, and were therefore not used. Interestingly, LMP2A KO or EBNA-LP KO EBV infected B cells were less sensitive to methionine restriction than cells infected by wildtype EBV or by the other EBV KO viruses (**Figure 5A**), suggesting their potential roles in metabolism remodeling.

**Figure 5.**
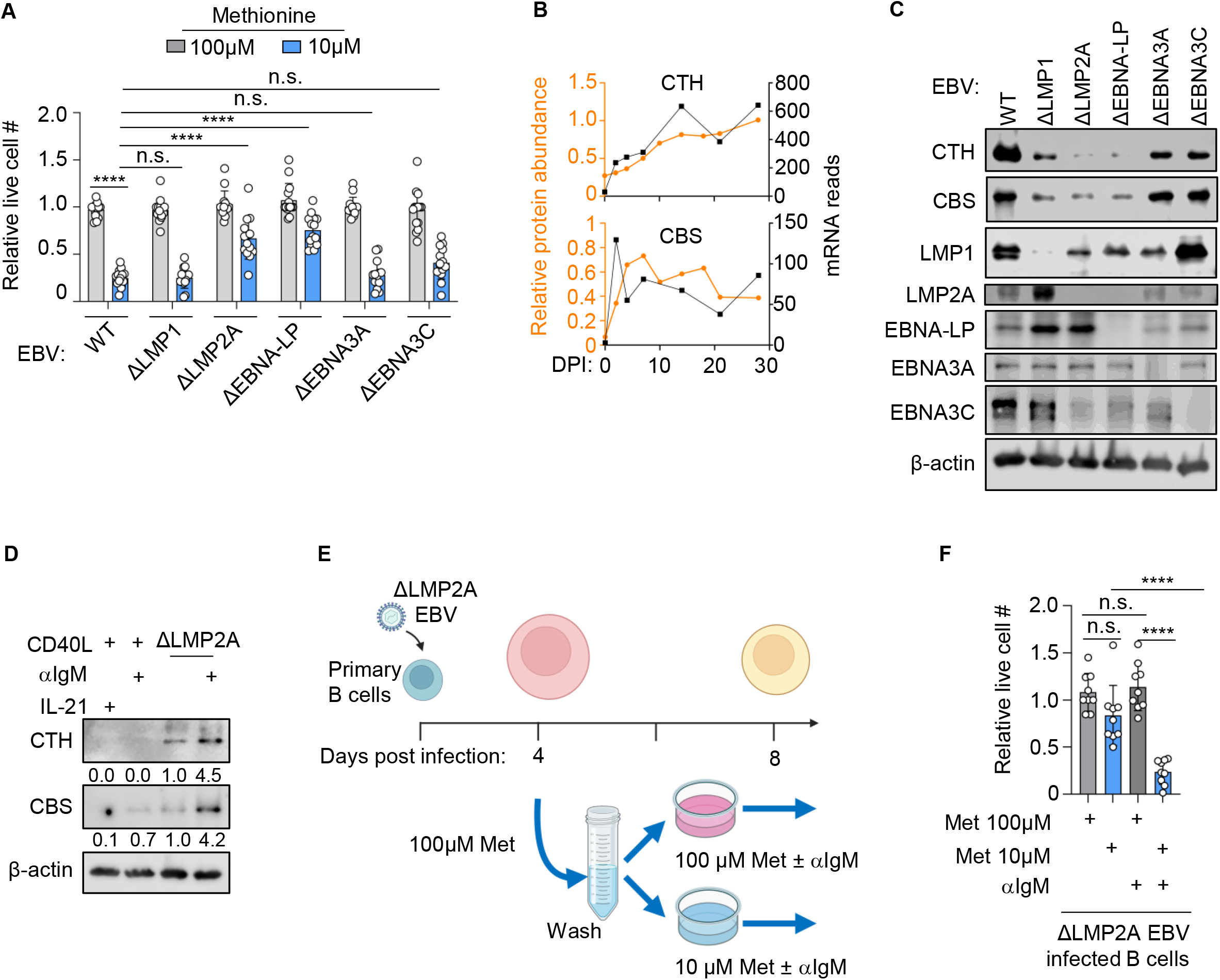
Latency III oncogene roles in EBV-induced B cell methionine dependency and transsulfuration pathway upregulation. (A) EBV oncogene effects on newly infected primary B cell methionine survival dependence. Mean ± SD relative live cell values from n=3 donors (3 replicates/donor) of peripheral blood B cells infected with EBV wildtype (WT) or knockout for the indicated oncogene^49^ at a multiplicity of infection of 0.3 for 2 days in methionine replete media. Cell numbers were normalized and cells were then cultured in media with 100 versus 10μM Met for 3 days. Mean cell number for each methionine replete condition were normalized to one. (B) EBV upregulation of transsulfuration pathway enzyme expression in newly infected primary B cells. Relative CTH and CBS protein abundances^8^ (left y-axis) versus mRNA levels^90^ (right y-axis) changes over the first 28 days of human peripheral blood B cell infection. (C) EBV oncogene effects on newly infected B cell transsulfuration pathway enzyme expression. Immunoblot analyses of WCL from peripheral blood B cells infected by the indicated EBV WT or KO strain and grown in methionine replete media for five days. Blot is representative of n=3 replicates. (D) B-cell receptor signaling rescues transsulfuration pathway enzyme expression in ΔLMP2A infected B cells. Immunoblot analyses of WCL from uninfected B cells stimulated by CD40 ligand (CD40L, 10ng/ml) and IL-21 (100ng/ml) or αIgM crosslinking (1μg/ml) for two days, or infected by LMP2A KO EBV for two days and then mock-stimulated or stimulated by αIgM crosslinking for two days. Cells were grown in methionine replete media. (E) Experimental scheme. Peripheral blood B cells were infected by ΔLMP2A EBV for 2 days and cultured in media with 100 or 10μM Met for four days. Media was refreshed, and cells were mock-stimulated or stimulated by αIgM crosslinking for four days. (F) B-cell receptor crosslinking effects on ΔLMP2A EBV infected B cell methionine survival dependence. Mean ± SD relative live cell numbers of peripheral blood B cells from n=3 donors (3 replicates each) infected by ΔLMP2A EBV and treated as in (E). One-way ANOVA was used for statistical tests in (A, F). ****p<0.0001, ns, non-significant.

Using our published proteomic dataset^8^, we identified that EBV rapidly induces both CBS and CTH expression in newly infected primary B-cells on the mRNA and protein levels (**Figure 5B**). We therefore next used the panel of recombinant EBV to explore specific EBV oncoprotein roles in transsulfuration enzyme expression. CBS and CTH were expressed in B cells infected by wildtype EBV at day 5 post-infection (**Figure 5C**). However, CBS and CTH expression were lower in cells infected by LMP2A KO, EBNA-LP KO, or to a lesser extent LMP1 KO EBV, suggesting that crosstalk between each of these viral oncogenes may upregulate the transsulfuration pathway. We note that LMP2A expression was also highly impaired in the EBNA-LP KO infected cells, consistent with a previous report ^55^. By contrast, EBNA-LP level increased with LMP2A KO (**Figure 5C**), suggesting that its expression is not sufficient to induce transsulfuration pathway enzyme expression. Taken together, our data suggest that LMP2A plays a central role in EBV-mediated CBS and CTS upregulation.

LMP2A mimics aspects of B-cell receptor immunoglobulin signaling to activate downstream pathways, including PI3K/mTORC1^24,56,57^. We therefore tested whether B-cell receptor stimulation could rescue CBS and CTH expression in peripheral blood B cells infected by LMP2A KO EBV. Primary B cells were infected by LMP2A KO EBV and grown in methionine replete media for two days. They were split into parallel cultures, which were either mock stimulated, or stimulated by αIgM crosslinking to drive B-cell receptor signaling for three additional days. For direct cross-comparison, primary B cells from the same donor were stimulated for three days by CD40 ligand (CD40L), whose signaling is mimicked by LMP1, and also by either the germinal center cytokine IL-21 or by αIgM to mimic physiological B-cell activation pathways. We observed modest CBS expression in CD40L/αIgM stimulated cells, but not appreciably in CD40L/IL-21 stimulated cells. Interestingly, αIgM stimulation strongly induced both CBS and CTH expression in LMP2A KO infected B-cells, suggesting that it compensated for LMP2A loss, and indicating that B-cell receptor pathway activation by either LMP2A or by immunoglobulin signaling can induce transsulfuration pathway enzyme expression in EBV-infected B cells (**Figure 5D**).

We next investigated whether B-cell receptor signaling sensitizes LMP2A KO EBV infected cells to methionine restriction driven cell death. Peripheral blood B cells were infected by LMP2A KO EBV and cultured for 4 days in media with 100μM methionine for four days. We then shifted cells into parallel cultures with media containing 100μM vs 10 μM methionine, in the absence or presence of αIgM stimulation (**Figure 5E**). Interestingly, whereas aIgM stimulation did not sensitize LMP2A KO infected B cells to cell death in methionine replete media, it triggered LMP2A KO infected B cell death when grown under methionine restriction (**Figure 5F**). By contrast, αIgM/CD40L co-stimulation, or CD40L/IL21 co-stimulation did not induce primary B cell death under methionine replete or restriction conditions (**Figure S5A-B**). These data suggest that in the context of EBV infection, LMP2A, or B cell receptor signaling, induce transsulfuration and sensitize cells to methionine restriction induced cell death.

### Dietary methionine restriction impairs LCL Xenograft outgrowth *in vivo*

Methionine is an essential amino acid, and dietary methionine restriction can therefore significantly reduce plasma methionine levels, including in murine models^58^. Based on our prior work, dietary methionine restriction downmodulates NOD *SCID* IL2 receptor gamma (NSG) mouse serum methionine concentration by ∼50% within one week ^29^. Given the above results, we hypothesized that the outgrowth of LCL xenografts, which model immunosuppressed host immunoblastic lymphomas such as PTLD, would be impaired by dietary methionine restriction.

To test this hypothesis, GM12878 LCL tumor xenografts were implanted subcutaneously in the flanks of 28 NSG mice (equal numbers of male/female, approximately 10-11 weeks of age)^59^. Tumors were allowed to grow for one week, at which point they were palpable. Mice were randomized to control methionine-replete (0.86% w/w methionine) versus methionine-restricted (0.086% w/w methionine) diets, which had balanced total amino acid levels (**Figure 6A**). Dietary methionine restriction impaired xenograft outgrowth across the time course: tumor volumes diverged from controls and by day 15 post-diet change, at which point they were reduced significantly (**Figure 6B**). This discrepancy continued to widen through day 28, where LCL xenograft tumor volumes were reduced by approximately 50% (methionine restricted, 961 mean ± SEM mm³; control, methionine replete 1781 mean ± SEM] mm³; n=4/group; p<0.0001). Body weights did not differ significantly between mice fed control or methionine restricted diet, indicating that dietary effects on tumor size were not attributable to overt systemic toxicity (**Figure S6A**).

**Figure 6.**
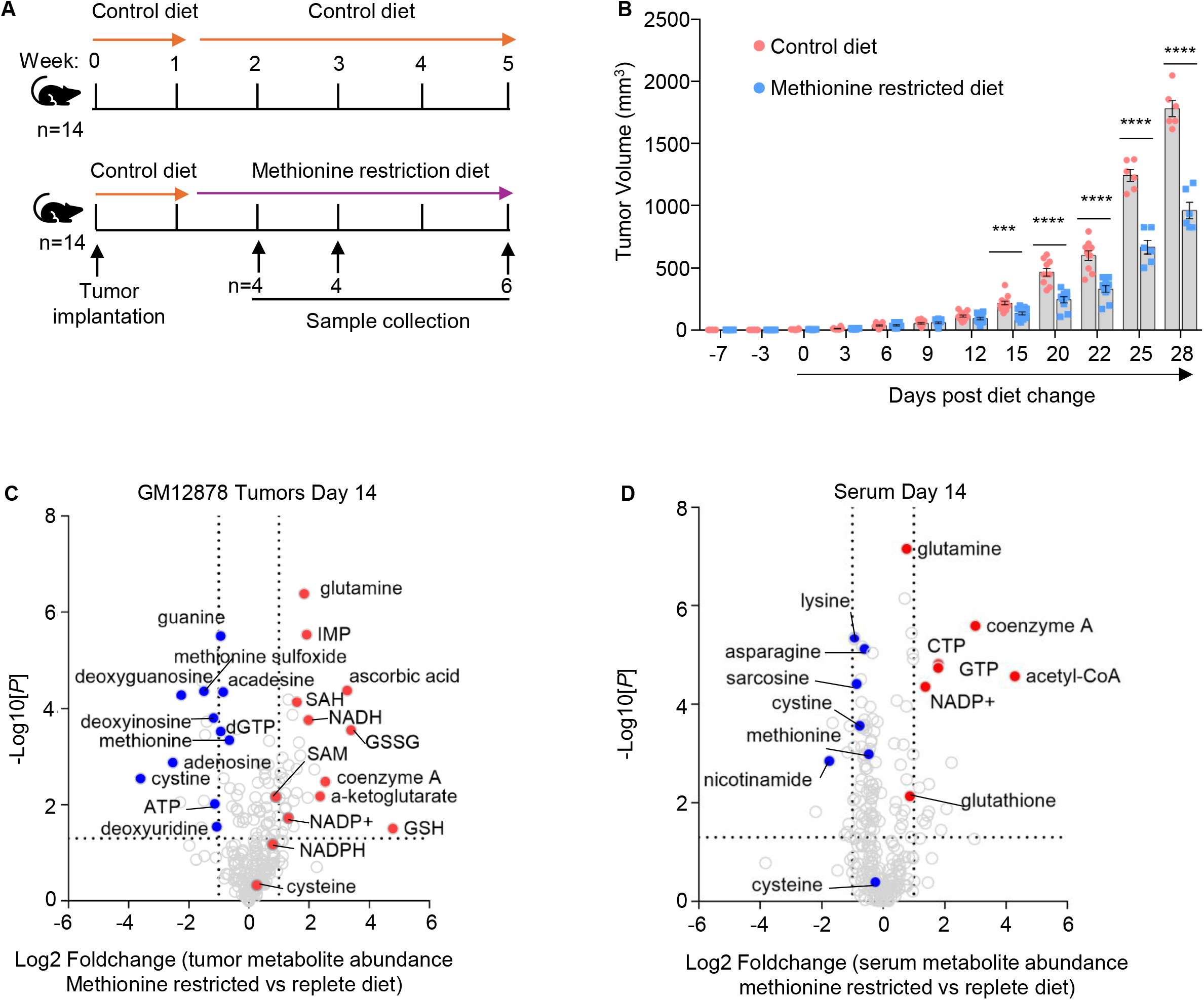
Dietary methionine restriction impedes LCL xenograft growth and strongly depletes tumor cystine in vivo. (A) Xenograft experimental scheme. GM12878 LCL tumors were established in NOD *scid* gamma mice flanks. Mice were fed a methionine replete control diet for 1 week. Mice were then randomized to control versus methionine restricted diets for four weeks. (B) Dietary methionine restriction effects on GM12878 tumor volumes. Mean ± SD LCL tumor volumes in mice fed control (red) versus methionine restricted (blue) diets. Student’s t-test, ***p<0.001, ****p<0.0001. (C) Dietary methionine restriction effects on GM12878 tumor metabolomes. Volcano plot from LC-MS polar metabolite analysis of n=4 LC-MS replicates of GM12878 xenografts harvested from mice fed control versus methionine restricted diets for 2 weeks. (D) Dietary methionine restriction effects on NOD-SCID serum metabolomes. Volcano plot from LC-MS polar metabolite analysis of n=4 LC-MS replicates of NOD-SCID gamma mice as in (C).

To gain further insights into how dietary methionine altered LCL xenografts, we performed metabolomic analyses of serum and of tumors explanted at day 14 of methionine replete versus restricted diets. As expected, dietary methionine restriction reduced xenograft tumor methionine levels by nearly 40%, as compared with a nearly 30% reduction in serum methionine levels (**Figure 6C-D, Table S5**). Interestingly, cystine was the most highly depleted metabolite in LCL tumors grown under dietary methionine restriction, reaching nearly 16-fold reduction, even though cysteine levels were not significantly changed (**Figure 6C, Table S5**). By contrast, dietary methionine restriction only reduced serum cystine concentration by nearly 2-fold (**Figure 6D**). This difference may likely be explained by the significantly higher dietary methionine restricted LCL xenograft levels of reduced (GSH) and oxidized (GSSG) glutathione, which were approximately 22-fold and 10-fold higher, respectively (**Figure 6C-D**). Dietary methionine restriction instead increased serum GSH by ∼2-fold, but decreased GSSG by a similar level (**Figure 6D, Table S5**). We therefore suspect that dietary methionine restriction increased cysteine incorporation into newly synthesized glutathione, potentially as a redox defense mechanism to increased lipid ROS. Consistent with *in vitro* methionine restriction, dietary methionine restriction mildly increased LCL xenograft tumor PARP1 cleavage (**Figure S6B**).

### Methioininase and cyst(e)inase treatment impedes LCL xenograft growth *in vivo*

Given our observation that dietary methionine restriction strongly reduced LCL xenograft cystine levels, we hypothesized that LCL xenograft growth would be highly sensitive to combined methionine and cyst(e)ine depletion. Notably, cysteine is a non-essential amino acid and can be safely depleted^60,61^. To test this hypothesis, we employed recombinant methionine and cyst(e)ine degrading enzymes, termed methioninase and cyst(e)inase, respectively ^60,62,63^. Both are derived from the human enzyme cystathionine-γ-lyase^60,63^. PEGylated methioninase and cyst(e)inase (cystathionine-γ-lyase) enzymes strongly deplete serum methionine and cyst(e)ine levels, respectively, are well-tolerated in mammals and have long half-lives.

GM12878 LCL tumor xenografts were implanted in flanks of 16 NSG mice (equal numbers male/female). Tumors were allowed to grow for 16 days to reach a size of 30-50mm^3 per tumor, at which time mice were randomized to receive vehicle, methioninase (90mg/kg), cyst(e)inase (1.5mg) or both via the intraperitoneal route. Mice were dosed for six doses (**Figure 7A**). Methioninase and/or cyst(e)inase were well tolerated and did not significantly alter mouse body weight (**Figure S7A**). Consistent with our dietary methionine restriction results, methioninase administration significantly reduced LCL xenograft outgrowth (**Figure 7B**).

**Figure 7.**
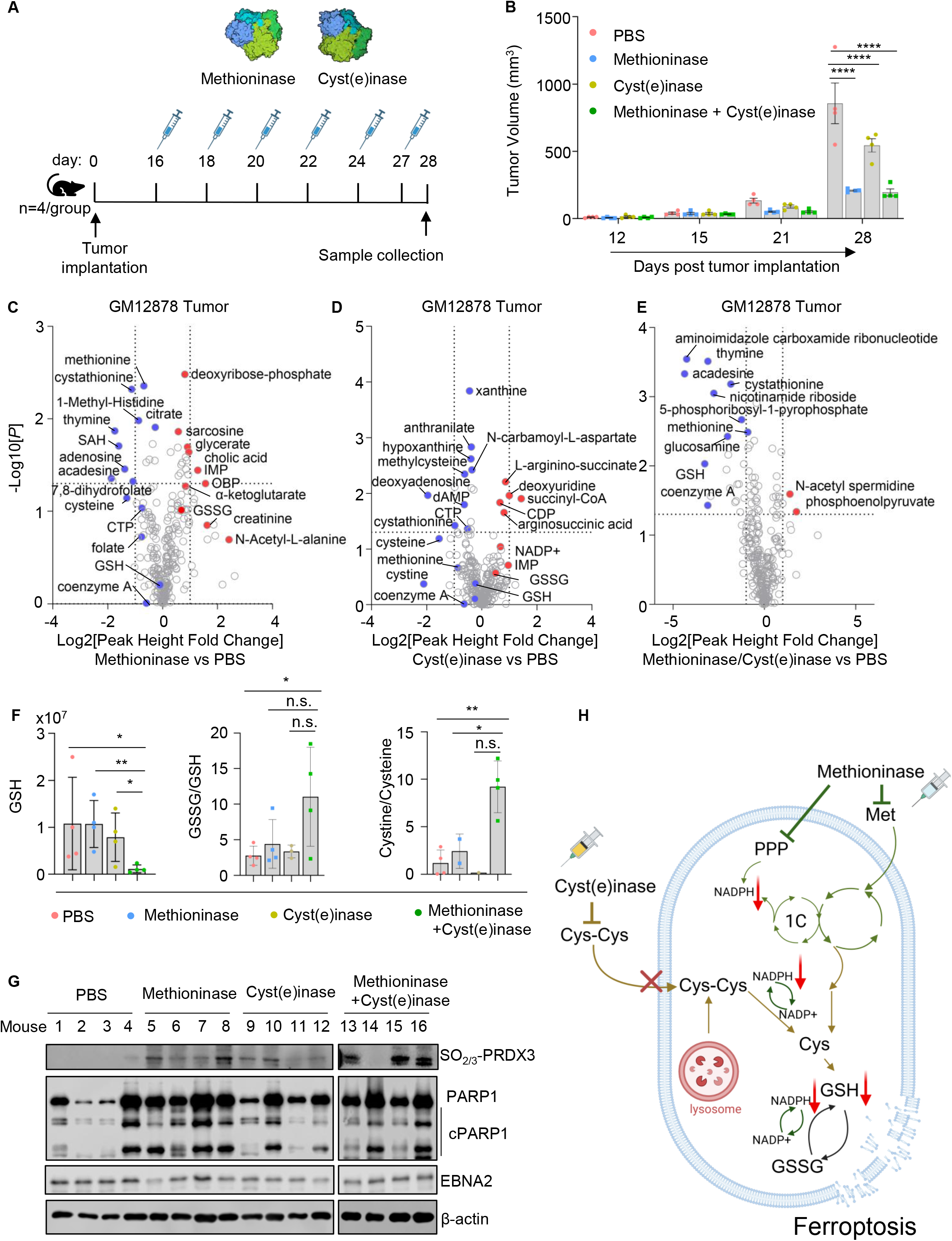
Methioninase and cyst(e)inase effects on GM12878 xenografts in vivo. (A) Xenograft experimental scheme. GM12878 LCL tumors were planted in NOD-*scid* gamma mouse flanks. Mice were administered six doses of PBS vehicle, methioninase (90mg/kg) or cyst(e)inase (1.5mg/mouse) intraperitoneally between days 16-27. (B) Methioninase and cyst(e)inase effects on GM12878 tumor volumes. Mean ± SD LCL tumor volumes in mice administered PBS vehicle (red), methioninase (blue) cyst(e)inase (yellow) or both (green). (C) Methioninase effects on GM12878 tumor metabolomes. Volcano plot from LC-MS polar metabolite analysis of n=4 LC-MS replicates of GM12878 xenografts explanted at day 28 from mice that were administered PBS versus methioninase. (D) Cyst(e)inase effects on GM12878 tumor metabolomes. Volcano plot from LC-MS polar metabolite analysis of n=4 LC-MS replicates of GM12878 xenografts explanted at day 28 from mice that were administered PBS versus cyst(e)inase. (E) Combined methioninase/cyst(e)inase effects on GM12878 tumor metabolomes. Volcano plot from LC-MS polar metabolite analysis of n=4 LC-MS replicates of GM12878 xenografts explanted at day 28 from mice that were administered PBS versus methioninase + cyst(e)inase. (F) Methioninase and cyst(e)inase effects on LCL tumor glutathione and cyst(e)ine abundances. Mean ± SD GSH, GSSG/GSH ratio or cystine/cysteine ratio ion intensities from LC-MS analyses shown in (C). (G) Methioninase and/or cyst(e)inase effects on LCL ferroptosis. Representative immunoblot analysis from n=3 replicates of WCL from Day 28 tumors. Hyperoxidized peroxiredoxin-3 (PRDX3-SO_2/3_), a marker of oxidative stress and ferroptosis. (H) Schematic model of methioninase + cyst(e)inase injection. Student’s t-test was used for statistical analyses in (A, F). ****p<0.0001, **p<0.01, *p<0.05. ns, non-significant.

Cyst(e)inase administration also reduced xenograft tumor size, albeit to a lesser extent than methioninase. Combined methioninase and cyst(e)inase shrunk tumors, to a somewhat greater extent than methioninase alone, where LCL xenograft tumor volumes were reduced by approximately 70% (PBS, 857 mean ± SEM mm³; methioninase, 210 mean ± SEM mm³, cyst(e)inase 545 mean ± SEM mm³, combined, 195 mean ± SEM mm³) (**Figure 7B**).

To gain further mechanistic insights into methioninase and cyst(e)inase effects, we performed LC/MS profiling of explanted tumor xenografts and serum at day 28. (**Figure 7A**, **C-E, S7B-E**). When dosed alone, methioninase reduced serum methionine concentration by 75%, whereas cyst(e)inase nearly completely depleted cystine. When dosed together, serum methionine and cystine levels were similarly reduced (**Figure S7C-E, Table S7**). Consistent with dietary methionine restriction, methioninase strongly depleted serum cystine levels (**Figure S7C**, **Table S7**).

Tumor LC/MS PCA of amino acid metabolites resolved the four experimental groups into largely discrete clusters, indicating distinct global metabolomic profiles across conditions (**Figure S7F**). Methioninase perturbed a range of metabolomic pathways, with cysteine and methionine metabolism amongst the most highly perturbed (**Figure S7F**). Methioninase reduced LCL tumor methionine levels by approximately 40%. Interestingly, cyst(e)inase treatment reduced tumor methionine by a similar extent, and combined methioninase/cyst(e)inase treatment further reduced tumor methionine (**Figure 7E, Table S6**). These data are consistent with a model in which serum cystine depletion forces LCL tumors to become increasingly reliant on transsulfuration to meet cysteine demand, including for glutathione synthesis. In support, transsulfuration pathway intermediate cystathionine levels were reduced to a similar extent (approximately 50%) by methioninase and cyst(e)inase treatment, whereas combined methioninase/cyst(e)inase treatment reduced cystathioinine by over nearly 75% (**Figure 7C-E, Table S6**).

Combined methioninase/cyst(e)inase treatment had intriguingly distinct effects on tumor cystine and cysteine levels than treatment with either alone (**Figure S7G-I**). Tumor cystine levels were reduced by nearly 2-fold by methioninase, likely reflecting methioninase effects on serum cystine levels, and were nearly completely depleted by cyst(e)inase. However, combined methioninase/cyst(e)inase treatment instead highly increased cystine levels (**Figure 7E, Table S6**). While treatment with methioninase and with cyst(e)inase likewise strongly decreased tumor cysteine levels, cysteine abundance was similar in tumors from vehicle control versus methioninase/cyst(e)inase treated mice (**Figure 7F, Table S6**).

Cysteine is the rate-limiting amino acid in glutathione synthesis. We therefore next inspected enzyme effects on tumor glutathione levels. GSH and GSSG levels were little changed by methioninase treatment (**Figure 7C, Table S6**), in contrast with dietary methionine restriction, which significantly increased both. We suspect that this difference is related to methioninase effects on serum cystine and cysteine, each of which were reduced by nearly 80% (**Figure S7C**). Cyst(e)inase decreased GSH levels by approximately 35%, likely reflecting the near total depletion of serum cysteine and supporting the hypothesis that LCLs can utilize transsulfuration to maintain tumor cysteine and glutathione levels (**Figure 7D**, **Table S7**). Xenograft GSSG levels were not significantly changed by cyst(e)inase administration.

Intriguingly, combined methioninase and cyst(e)inase administration strongly reduced LCL xenograft tumor GSH levels by nearly 90% and also increased GSSG by nearly 40% (**Figure 7E-F, Table S6**). This likely occurred because combined enzyme administration depleted both serum methionine and cystine, limiting the ability of tumor cells to compensate for loss of cystine uptake by transsulfuration mediated *de novo* cysteine biosynthesis. Furthermore, combined methioninase and cyst(e)inase administration reduced NADPH levels by ∼33% and also decreased the NADPH/NADP ratio (**Figure 7E, Table S6**). We note that combined enzyme administration highly perturbed both the pentose phosphate pathway and one carbon and folate metabolism, each of which supply NAPDH in EBV-transformed B-cells^8^ (**Figure S7H**). Combined enzyme administration also highly altered the pantothenate and CoA biosynthesis pathway, including by strongly reducing CoA and pantothenate levels (**Figure S7H, Table S6**). This contrasted with dietary methionine restriction, which instead increased serum and tumor CoA levels.

To then investigate methioninase and/or cyst(e)inase effects on xenograft cell death pathways, we performed immunoblot analysis of whole cell extracts prepared from day 28 tumors. We blotted for PAPR cleavage as a readout of caspase activity and for hyperoxidized peroxiredoxin 3 (PRDX3) as a readout of ferroptosis ^64^. Whereas methioninase or cyst(e)inase alone induced hyperoxidized peroxiredoxin 3 (PRDX3), combined treatment did so more robustly, suggesting a higher degree of ferroptosis induction (**Figure 7G**). PARP cleavage was observed in each enzyme treated group, though appeared most robust with methioninase treatment alone (**Figure 7G**). Taken together, these data suggest that combined methioninase/cyst(e)inase administration blocked the ability of LCL tumors to fill GSH pools, likely both due to depletion of intracellular cysteine through impaired transsulfuration, loss of cystine uptake, and through reduction in NAPDH, causing GSH depletion and ferroptosis induction (**Figure 7H**).

## DISCUSSION

Tumor cells frequently rewire methionine metabolism, and most acquire a dependence on exogenous methionine for proliferation even with homocysteine supplementation, which is termed the Hoffman effect^40,65^. Here, we found that EBV-transformed LCLs are not merely methionine-dependent for growth, but are critically dependent on methionine for survival. Methionine restriction rapidly remodeled the LCL transcriptome and metabolome, elevated lipid ROS, and triggered ferroptosis. Iron chelation, cysteine supplementation, or lipid-ROS scavenging rescued LCL survival, whereas pan-caspase or necroptosis blockade did not. Metabolic tracing localized the underlying vulnerability to the transsulfuration pathway, which the transforming latency III program strongly induced to supply cysteine for glutathione synthesis and NADPH-dependent redox defense. The EBV oncoprotein LMP2A supported transsulfuration enzyme expression and methionine dependence in newly infected primary human B cells, and dietary methionine restriction, more potently combined with cyst(e)inase, depleted LCL xenograft glutathione and drove ferroptosis *in vivo*.

That LCLs undergo ferroptosis rather than a survivable arrest upon methionine withdrawal is highly unusual among cancer cells and reframes their methionine dependence as fundamentally a redox-survival dependence. Lymphoblastoid cells become progressively ferroptosis-resistant as they undergo EBV-mediated immortalization, but LCLs nonetheless rely on GPX4- and glutathione-mediated detoxification of lipid ROS^37^. Our data indicate that methionine restriction re-opens this latent vulnerability by severing a key cysteine supply line. This adds EBV-transformed B cells to a small but growing set of contexts in which methionine flux, acting through transsulfuration-derived cysteine, alters the ferroptosis threshold^18,66–69^. It suggests that the LCL redox program is unusually reliant on methionine-derived, rather than solely imported cysteine.

Our findings suggest that the methionine vulnerabilities of latency I versus latency III B cells are mechanistically distinct. In EBV+ Burkitt cells, which maintain a heavily methylated genome and the restricted latency I program^70–72^, methionine restriction lowers the SAM/SAH ratio, hypomethylates the viral episome, and de-represses EBV latency and lytic genes^29,30^. In LCLs, by contrast, methionine restriction did not significantly change the SAM/SAH ratio or broadly alter EBV transcript abundances, and instead precipitated a redox catastrophe. EBV therefore appears to impose two separable methionine dependencies across its lifecycle: an epigenetic dependence that enforces restricted latency in Burkitt and possibly also memory B cells, which share the latency I program, and a redox/survival dependence in proliferating latency III cells. These have opposite therapeutic logics, since methionine restriction de-represses potentially immunogenic viral antigens in latency I tumors, but is directly cytotoxic to latency III cells.

Mechanistically, LMP2A emerged as a central driver of transsulfuration and of methionine dependence, with contributions from EBNA-LP and LMP1. Notably, LMP1 and LMP2A are often co-expressed and co-localize at membrane signaling sites^73^. We recently reported that LMP1 signaling supports LCL erastin resistance through upregulation of PFKFB4, PPP and NADPH pools^74^. Because EBNA-LP loss also impaired LMP2A expression, its effect on CBS and CTH may be at least partly indirect. Immunoglobulin crosslinking phenocopied LMP2A, inducing CBS and CTH in LMP2A-knockout-infected cells and restoring methionine-restriction sensitivity, indicating that B-cell receptor pathway activation is sufficient to upregulate transsulfuration in EBV-infected B cells. LMP2A and the B cell receptor (BCR) both signal through PI3K/mTORC1^24^, and PI3K-driven transsulfuration is expected to consume homocysteine for cysteine synthesis at the expense of its remethylation to methionine^75^. This diversion would simultaneously support redox defense and deepen the requirement for exogenous methionine. This positions transsulfuration as an oncogene-tunable node linking BCR-mimicry to redox homeostasis, and raises the possibility that physiological germinal-center BCR signals similarly calibrate B-cell transsulfuration and ferroptosis sensitivity.

We previously reported that methionine restriction suppresses expression of multiple EBV oncogenes in newly infected cells^29^. An unexpected observation was that Fer-1 and NAC not only rescued newly infected B-cell survival under methionine restriction, but also restored EBNA2 and LMP1 expression, shifting cells from death toward a Hoffman-effect-like arrest. This suggests that methionine-restriction-induced lipid ROS suppresses EBV oncogene expression, and that redox stress may both kill and transcriptionally disarm transforming cells. Whether this reflects oxidative effects on oncogene transcription, translation, or protein stability remains to be defined, but such a feed-forward loop could sharpen the sensitivity of newly infected cells to methionine limitation during the earliest, most vulnerable phases of transformation. Given that EBNA2 and LMP1 are increasingly implicated in the pathogenesis of multiple sclerosis and systemic lupus erythematosus, it will be of interest to determine methionine restriction or depletion effects, including on the EBV+ CD27+CD21^low^ memory cell reservoir^7,76–78^.

These findings have translational implications for EBV+ latency III malignancies, in particular PTLD and immunoblastic diffuse large B-cell lymphoma. Notably, dietary methionine restriction paradoxically increased tumor glutathione, an adaptive redox response likely supported by residual serum cystine, which may limit single-agent efficacy. Combined methioninase and cyst(e)inase administration overcame this compensation: by depleting both extracellular methionine and cystine, it blocked cystine import and transsulfuration-mediated *de novo* cysteine synthesis in parallel, collapsing tumor glutathione by ∼90%, raising GSSG, lowering NADPH and CoA, and inducing ferroptosis as marked by hyperoxidized PRDX3. Dual amino-acid-depleting enzyme therapy therefore represents a rational strategy to force ferroptosis in EBV-transformed B cells.

Our study has limitations. Xenografts in immunodeficient mice omit adaptive immune contributions, yet methionine restriction also enhances anti-tumor CD8+ T-cell function, and the innate-immune (OAS/ISG) signature we observed under methionine restriction hints at additional host-directed effects to be explored in immunocompetent models. Whether disulfidptosis^79,80^ contributes under the high-SLC7A11, cystine-loaded conditions produced by combined enzyme treatment, and whether primary PTLD share this transsulfuration dependence, remain important open questions.

In summary, our results suggest that EBV-infected lymphoblastoid B cells are highly dependent on extracellular methionine for survival. Methionine restriction triggers LCL lipid ROS production and rapid cell death. LMP2A expression was critical for such methionine dependence, likely through PI3K pathway activation. Methionine depletion strongly impaired LCL xenograft outgrowth *in vivo*. Combined cyst(e)inase/methioninase administration depleted LCL xenograft glutathione and induced ferroptosis.

## Supporting information

Table S1. Methionine Restiction effects on the LCL transcriptome in vitro

Table S2. Methionine restriction effects on the LCL metabolome in vitro

Table S3. Methioinine tracing analyses in GM12878, MUTU I and III

Table S4. Metabolomic analyses of MUTU I vs III

Table S5. Dietary methioinine restriction effects on GM12878 xenografts and serum

Table S6. Methioininase and Cyst(e)inase effects on NSG GM12878 xenografts

Table S7. Methioininase and Cyst(e)inase effects on NSG serum

Reagents and Resources

## ACKNOWLEDGMENTS

This work was supported by American Cancer Society Post-doctoral Fellowships PF-24-1308318-01-TBE to S.W and PF-24-1250090-01-IBCD to BM, by R00DE031016 to R.G., and by R01DE033907, R01CA228700, U01CA275301 and P01CA269043 to B.E.G, and P01CA120964 to JMA. S.F.L was supported by NIH P30 AI060354 Harvard University Center for AIDS 508 Research (CFAR), which is supported by the following NIH Co-Funding and 509 Participating Institutes and Centers: NIAID, NCI, NICHD, NHLBI, NIDA, NIMH, NIA, 510 NIDDK, NIMHD, NIDCR, NINR, OAR, and FIC. The authors also acknowledge generous philanthropic support of George and Sandra K. Schussel. G.G. and E.S. have an equity interest in Aeglea Biotherapeutics, a company that has licensed the commercial development of cyst(e)inase, and are inventors on U.S. patents and corresponding applications and patents in other countries, all submitted by the University of Texas, related to l-cyst(e)inase: 10,363,311 and 20180327734. The other authors declare no competing interests, financial or otherwise. The funders had no role in study design, data collection and interpretation, or the decision to submit the work for publication. The authors acknowledge Dr. Wolfgang Hammerschmidt for generously sharing EBV B95.8 wildtype and oncogene knockout producer cells.

## AUTHOR CONTRIBUTIONS

R.G. and B.E.G. conceived the initial project direction. R.G. and B.E.G. supervised the study. R.G. made the original observation that EBV-transformed LCLs are selectively sensitive to methionine restriction. R.G. generated several core datasets that formed the foundation of the study, including RNA-seq, stable-isotope tracing, dietary methionine-restriction mouse studies, and MR tumor metabolomics. S.W. completed the study by performing additional mechanistic experiments, developing the final ferroptosis story. B.M. and H.L. produced the recombinant EBV viruses. Y.L. and S.F.L. performed immunoblot experiments. R.P. runs an animal core facility and executed the mouse xenograft experiment. J.M.A. runs metabolomics core facility and performed LC/MS analysis. R.G., E.S., G.G. and B.E.G. designed the methioniase/cysteinase in vivo experiments. E.S. and G.G. produced the Methioninase and Cyst(e)inase. S.W. and R.G. performed bioinformatic analyses. S.W. and B.E.G. wrote the first draft of the manuscript. All authors analyzed the results, discussed the findings, reviewed the manuscript, and approved the final version.

## RESOURCE AVAILABILITY

Further information and requests for resources and reagents should be directed to and will be fulfilled by the lead contact, Benjamin Gewurz.

## Materials availability

All plasmids generated in this study will be made available on request

## Data and code availability

All RNA-seq datasets have been deposited to the NIH GEO omnibus. The accession number for the RNA-seq dataset reported in this paper is GEO GSE336950.

Figures were drawn with commercially available GraphPad, Biorender, Microsoft Powerpoint. This study did not generate any original code.

## EXPERIMENTAL MODEL AND SUBJECT DETAILS

### Cell lines and reagents

The EBV+ Burkitt cell lines P3HR-1, Akata, Mutu I, MUTU III and Raji were obtained from Drs. Jeffrey Sample or Elliott Kieff. B95.8 producer cells were obtained from Elliott Kieff. GM12878 and GM13111 LCLs were obtained from Coriell. Cells were cultured in a humidified incubator at 37°C with 5% CO2 and routinely tested and certified as mycoplasma-free using the MycoAlert kit (Lonza). B cells were grown in RPMI 1640 medium (Gibco, Life Technologies) with 10% fetal bovine serum (FBS, Gibco) or with dialyzed FBS where indicated (Gibco). HEK-293T were obtained from ATCC and grown in Dulbecco’s Modified Eagle’s Medium (DMEM) with 10% FBS. For selection of transduced cells, puromycin was added at the concentration of 3 μg/mL. EBV⁺ Akata cells were obtained from Elliott Kieff. B95.8 cells with 4-Hydroxytamoxifen(4-HT)-inducible lytic cycle was obtained from Dr. Eric Johannsen.

Deferoxamine mesylate (DFO), an iron chelator, was obtained from Sigma and used at 100nM. N-acetyl-L-cysteine (NAC), an antioxidant and ROS scavenger, was obtained from Millipore Sigma and used at 10mM. Z-VAD-FMK (benzyloxycarbonyl-Val-Ala-Asp(OMe) fluoromethylketone), a pan-caspase inhibitor, was obtained from SelleckChem and used at 20 μM. Necrostatin-1 (Nec-1), a RIPK1/necroptosis inhibitor, was obtained from Apexbio and used at 160μM. DL-Homocysteine thiolactone hydrochloride (Hcy) was obtained from Thermo Fisher and used at 100μM. Ferrostatin-1 (Fer-1) was obtained from SelleckChem and used at 2.5μM. Erastin was obtained from SelleckChem and used at 10μM. Cysteamine was obtained from Sigma and used at 100μM. Cystathionine γ-lyase (CTH) inhibitors β-cyano-L-alanine and DL-propargyl glycine were both obtained from Cayman and used at 100μM and 1mM respectively.

### Primary human B cells

De-identified leukocyte fractions remaining from platelet donations were obtained from the Brigham and Women’s Hospital Blood Bank. All discarded blood cells were collected following platelet donation, in accordance with institutional guidelines. Donor gender and age information was not available due to the donor de-identification. Research involving primary human blood cells was reviewed and approved by the Brigham and Women’s Hospital Institutional Review Board, protocol 2022p001270. Primary human B cells were isolated by negative selection using the RosetteSep Human B Cell Enrichment and EasySep Human B Cell Enrichment kits (Stem Cell Technologies), following the manufacturers’ instructions. The purity of isolated B cells was verified by detecting CD19 expression on the plasma membrane via FACS analysis. Cells were maintained in RPMI 1640 medium supplemented with 10% FBS, and freshly isolated primary B cells were plated in either control or MR media at a density of 1 × 10L cells per mL.

### Mice

NOD.Cg-*Prkdc*^scid^ *Il2rg*^tm1Wjl^/SzJ (NSG) mice were originally procured from Jackson Labs, strain #005557, and bred in house at Cornell University, Ithaca under the supervision of Cornell’s animal breeding program. 10-11 weeks old NSG mice (equal numbers of male and female) were used for this study under the approval of the IACUC committee (protocol #2017-0035). The Center for Animal Resources and Education (CARE) staff, a team of veterinarians and technicians, monitored animal health daily for abnormal behavior, body condition, and tumor burden. Animals were housed in microisolator cages, no more than five mice per cage, on a 12 h light/12 h dark light cycle, given water *ad libitum*, and food was given according to experimental design.

## METHOD DETAILS

### Live cell number quantitation

Cell cultures were mixed thoroughly, and 100μl cell culture was transferred from each biological replicate into U-bottom 96-well plates, pelleted at 200xg for 5min, and resuspended in 100μl PBS. 100μl of 0.4% Trypan Blue Solution (Thermo) was added and mixed. Live cells were counted on dual-chamber cell counting slides (BioRad) by a TC20™ automated cell counter (BioRad) with default settings.

### Flow cytometry analysis

Cells were washed once with PBS before staining. For 7-AAD and annexin V FACS analysis, 1×10^6^ cells were stained in 100μl of staining buffer that contained FITC-conjugated anti-annexin V antibody (1:20) and 5μg 7-AAD for 15 min. Annexin V antibody and 7-AAD were diluted in annexin V binding buffer, which contains 10 mM HEPES pH 7.4, 140 mM NaCl, and 2.5 mM CaCl_2_. Before flow cytometry analysis, the total volume was expanded to 500μl with annexin V binding buffer. For BODIPY (lipid ROS) analysis, 6×10^5^ cell were seeded in 200μl RPMI media containing 2.5μM BODIPY™ 581/591 C11 and incubated at 37°C for 30min. For H2DCFDA analysis, 6×10^5^ cells were seeded in 200μl RPMI media containing 1 μM H2DCFDA and incubated at 37°C for 30min. Cells were then washed twice with PBS containing 2% FBS and resuspended in 500μl PBS containing 2% FBS before FACS analysis. For cell cycle analysis, EdU (MedChemExpress) was diluted to 10 μM in RPMI 1640 culture media of cells and incubated for 30 min. Then cells were washed with PBS and fixed in 70% ethanol overnight. Fixed cell were incubated in a PBS buffer (Fisher Scientific) containing 20 μg/ml propidium iodide, 200μg/ml RNase A (DNase-and protease-free, Thermo Fisher) and 0.1% Triton-X (Sigma) at 37°C for 30 min. Cells were washed and incubated in PBS supplemented with 2mM CuSO4, 10mM sodium ascorbate (Sigma) and 25μg/ml Alexa Fluor™ 488 Azide (1:1000; Thermo Fisher) for 2 hrs at 37°C. FACS was performed on a BD FACSCalibur instrument or a Cytek Aurora and analysis was performed with FlowJo V10 software.

### Caspase-Glo 3/7 assay

Caspase-Glo 3/7 (Promega) was added to equal volume of cells in PBS, mixed and incubated for 30 minutes according to the manufacturer’s instructions, followed by luminescence measurements on a SpectraMax iD5e plate reader (Molecular Devices). Readings were divided by the total cell density determined by TC20™ automated cell counter (BioRad).

### Immunoblot analysis

Whole cell lysate (WCL) were mixed with Laemmli loading buffer, separated by SDS-PAGE, transferred onto nitrocellulose filters (Bio-Rad), blocked with TBST (Tris-buffered saline with Tween 20) containing 5% milk or 3% BSA (NEB), and then probed with primary antibodies at 4°C overnight, followed by secondary antibody (Cell Signaling Technology, catalog no. 7074, catalog no. 7076, catalog no. 7077; LI-COR INC, catalog no. 926-32210, catalog no. 926-32219) incubation for 1 h at room temperature. For all horseradish peroxidase (HRP)-conjugated secondary antibodies, ECL chemiluminescence (Millipore, catalog no. WBLUF0500) was used to develop HRP signal. All images were captured with a Li-Cor Fc platform. All antibodies used in this study are listed in reagent and resource table.

### RNAseq analysis

Total RNA was isolated by the RNeasy Mini kit (Qiagen), following the manufacturer’s manual. An in-column DNA digestion step was included to remove the residual genomic DNA contamination. To construct indexed libraries, 1 μg of total RNA was used for polyA mRNA-selection, using the NEBNext Poly(A) mRNA Magnetic Isolation Module (New England Biolabs), followed by library construction via the NEBNext Ultra RNA Library Prep Kit (New England Biolabs). Each experimental treatment was performed in triplicate. Libraries were multi-indexed, pooled and sequenced on an Illumina NextSeq 500 sequencer using single-end 75 bp reads (Illunima) at the Dana Farber Molecular Biology core. Adaptor-trimmed Illumina reads for each individual library were mapped back to the human GRCh37.83 transcriptome assembly (accession #: GCF_000001405) or EBV Akata genome (accession #: KC207813.1) using STAR2.5.2b^81^. Feature Counts were used to estimate the number of reads mapped to each contig^82^. Only transcripts with at least 5 cumulative mapping counts were used in this analysis. DESeq2 was used to evaluate differential expression (DE) ^83^. DESeq2 uses a negative binomial distribution to account for overdispersion in transcriptome datasets. It uses a conservative analysis that relies on a heuristic approach. Each DE analysis used pairwise comparison between the experimental and control groups. Differentially expressed genes were identified and a p values < 0.05 and absolute fold change > 2 cutoff was used. Differentially expressed genes were subjected to Enrichr analysis which was employed to perform gene list-based gene set enrichment analysis on the selected gene subset. The algorithm used to calculate combined scores was described previously^84^. P-value and log2 fold change were generated with DESeq2 under default settings with Wald test and normal shrinkage, respectively. Top 10 Enrichr terms that passed the adjusted p-value cutoff were visualized using Graphpad Prism 11. Volcano plots were built with Graphpad Prism11.

### EBV virus production, titration and infection of primary B cells

For Akata virus production, EBV⁺ Akata cells were resuspended in serum-free RPMI 1640 at a density of 2–3 × 10L cells/mL and induced for lytic reactivation with 10μg/mL goat anti-human IgG (Dako, Cat# A042402-2) for 72 h at 37 °C. For B95.8 virus production, B95.8 producer cells with 4-Hydroxytamoxifen(4-HT)-inducible lytic cycle^85^ were seeded in 10% FBS RPMI 1640 media containing 400nM 4-HT (Sigma) for 24hrs, followed by media change to 10% FBS RPMI1640 for additional 4 days. B95.8 virus titers were determined by percentage of cells expressing CD23 48 hours post-infection. For production of B95.8 BAC wildtype and mutant viruses^49^,293 producer cells, which were generously provided by Dr. Wolfgang Hammerschmidt, were reverse transfected at a seeding density of seeded at 4×10^6^ 5.5X10^6^/plate in a 100mm tissue culture plate. 6μg of the vector 509 MacVec-BZLF1 (EBV BZLF1 cDNA expression vector) and 6μg of the vector 2670_pRA-BZLF4 (EBV BALF4 cDNA expression vector) were pre-mixed with 72μl of 1mg/ml PEI MAX (Polysciences #24765-1) in 2.5ml Opti-MEM media (Thermo #31985070) and incubated for 15 minutes at RT before used to reverse transfect each 100mm plate of 293 BAC virus producing cells. The media was changed to RPMI 1640 supplemented with 10% fetal bovine serum 18 hours after transfection. 5-6 days after transfection, culture supernatants were collected, centrifuged at 1200rpm and 4000rpm sequentially, and further clarified by passage through a 0.45μm filter. Virions were pelleted by ultracentrifugation at 50,000×g (25,000 rpm) for 2.5 hours at 4C. Pellets were resuspended in PBS containing 2% dialyzed FBS, RPMI 1640 media, aliquoted, and stored at −80 °C until use. Viral stocks were thawed immediately before infection, and titers were determined by percentage of GFP+ cells on Raji B- cells.

For primary B cell transformation, freshly isolated primary human B cells described above were plated in 6-well plates at 1 × 106 cells/mL and EBV viruses were added to cells at an MOI=0.3. This MOI was chosen as an optimal dose for EBV-driven B cell outgrowth^49^. On the indicated days, cells were washed with PBS three times and then seeded in methionine-free media (Thermo Fisher). Metabolites, small molecules, αIgM antibody or cytokines were added as indicated in figure legends.

### Intracellular and serum metabolite profiling

GM12878 were washed with PBS three times and then resuspended in methionine free RPMI-1640 (ThermoFisher Cat # A1451701) supplemented with 10% dialyzed FBS (ThermoFisher Cat #26400044). L-methionine was added to the cell culture to 100μM or 1μM methionine. 3×10^6^ live cells were collected for intracellular metabolite profiling at 24 hours, a time point prior to cell death. Cells were pelleted at 200xg in falcon tubes and washed with 5 mL of room temperature PBS. Pellets were resuspended in 1 mL of dry ice cold 80% methanol, incubated at −80°C for 30 minutes and centrifuged at 21,000×g for 5 minutes to precipitate proteins. For mice xenograft tumors, 100mg of tumor tissues were washed once with PBC and submerged in 500 μl of 80% (vol/vol) HPLC-grade methanol (cooled to −80 °C). Tissues were ground in the tube for 1–2 min with small pestle/tissue grinder on dry ice, vortexed for 1 min at 4°C and incubated for 4 hours at −80 °C. Tubes were centrifuged at 21,000g for 10 min at 4°C to pellet insoluble proteins. For mice serum, 80 μl of HPLC-grade methanol was added to 20 μl of serum and incubated for 4 hours at −80 °C. Then tubes were centrifuged at 21,000g for 10 min at 4°C to pellet insoluble proteins.

Supernatants from above were collected in pre-chilled tubes and stored at −80°C. On the day of analysis, supernatants were incubated on ice for 20 minutes and clarified by centrifugation at 21,000×g at 4°C. At the Beth Israel Deaconess Mass Spectrometry core, supernatants were dried down in a speed vacuum concentrator (Savant SPD 1010, Thermofisher Scientific) and re-suspended in 100μL of 60/40 acetonitrile/water. The samples were then vortexed, sonicated in ice-cold water for 1 minute, and incubated on ice for 20 minutes. Supernatants were collected in an autosampler vial after centrifugation at 21,000 × g for 20 minutes at 4°C. Pooled Q-C samples were generated by combining ∼15μL of each sample.

Metabolite profiling was performed using Dionex Ultimate 3000 UHPLC system coupled to QExactive plus orbitrap mass spectrometer (ThermoFisher Scientific, Waltham, MA) with an Ion Max source and HESI II probe operating in switch polarity mode. A zwitterionic Sequent zic philic column (150 × 2.1mm, 5μm polymer, part # 150460, MilliporeSigma, Burlington, MA) was used for polar metabolite separation. Mobile phase A (MPA) was 20mM ammonium carbonate in water, pH9.6 (adjusted with ammonium hydroxide) and MPB was acetonitrile. The column was held at 27°C, injection volume 5μL, autosampler temperature 4°C and LC conditions at flow rate of 0.15 mL/min were: 0min: 80% B, 0.5min: 80% B, 20.5min: 20% B, 21.3min: 20%B, 21.5min: 80% B with 7.5min of column equilibration time. MS parameters were: sheath gas flow 30, aux gas flow 7, sweep gas flow 2, spray voltage 2.80kV for negative & 3.80kV for positive, capillary temperature 310°C, S-lens RF level 50 and aux gas heater temp 370°C. Data acquisition was done using Xcalibur 4.1 (ThermoFisher Scientific) and performed in full scan mode with a range of 70–1000m/z, resolution 70,000, AGC target 1e6 and maximum injection time of 80ms. Data analysis was performed in Compound Discoverer 3.1 and Tracefinder 4.1. Samples were injected in randomized order and pooled QC samples were injected regularly throughout the analytical batch. Metabolite annotation was done base on accurate mass (±5ppm) and matching retention time (±0.5min) as well as MS/MS fragmentation pattern from the pooled QC samples against in-house retention time +MSMS library of reference chemical standards. Metabolites with CV<30% in pooled QC were used for the statistical analysis. The quality of integration for each metabolite peak was reviewed. Metabolites with p-values < 0.05, log2(fold change)>1 or <−1 were used for pathway analysis using MetaboAnalyst 5.0 (https://www.metaboanalyst.ca/MetaboAnalyst/ModuleView.xhtml).

### Polar metabolite extraction and LC-MS/MS profiling

Prior to analysis, metabolite pellets were resuspended in 22 µL HPLC-grade water. Metabolomics data were acquired by liquid chromatography–tandem mass spectrometry (LC-MS/MS) as previously described^86^ at the Beth Israel Deaconess Medical Center Mass Spectrometry Facility. Briefly, 8 µL of each resuspended sample was injected and analyzed on a 6500 QTRAP hybrid triple quadrupole mass spectrometer (SCIEX) coupled to a Prominence UFLC HPLC system (Shimadzu) for steady-state metabolite profiling. Selected reaction monitoring (SRM) was used to quantify approximately 310 polar metabolites, employing positive/negative ion polarity switching with hydrophilic interaction liquid chromatography (HILIC) (Amide XBridge column, Waters). Q3 peak areas for each metabolite SRM transition were integrated using MultiQuant v3.2 software (SCIEX) and obtained results were normalized to total protein abundance as determined by BCA assay.

### ^13^C_5_-methionine isotopologue tracing

GM12878, MUTU I, or MUTU III were seeded at 3 × 10^5^/ml in methionine-free RPMI 1640 supplemented with 10% dialyzed FBS and 100 μM L-methionine (¹³C_5_, 99%; ¹LN, 99%) and cultured for 6 hours. 3 × 10^6^ cells were pelleted in falcon tubes at 200xg at 4°C for 5 min and washed with 5 mL of cold PBS. Pellets were resuspended in 1 mL of dry ice cold 80% methanol, incubated at −80°C for 30 minutes and centrifuged at 21,000 × g for 5 minutes to precipitate proteins. The supernatant was dried down in a speed vacuum concentrator at the Beth Israel Deaconess Mass Spectrometry core as described above. Intracellular metabolites were extracted and prepared for mass spectrometric analysis as described above. Q1/Q3 SRM transitions for ¹³C incorporation into polar metabolite isotopomers were established, and data were acquired by LC-MS/MS via SRM^87^. Peak areas were generated for each detected isotopomer using MultiQuant (version 3.2) software.

### *In vitro* methionine or cystine restriction

For methionine restriction, cells were washed with PBS three times and then resuspended in methionine free RPMI-1640 (ThermoFisher Cat # A1451701) supplemented with 10% dialyzed FBS (ThermoFisher Cat #26400044). L-methionine was added to the cell culture to the indicated concentrations and cells were grown for the indicated times, typically 30 hrs. For cystine restriction, cells were also washed with PBS three times and resuspended in replete or cystine-restricted media. Media for cystine restriction was made as described^88^. Briefly, RPMI 1640 medium without amino acids, sodium phosphate (Thermo Fisher) was reconstituted from power according to the manufacturer instruction, except that pH was not adjusted with HCl. Cystine stock solution was made by dissolving L-cystine 2HCl in 1N HCl to 50mg/ml or 0.5mg/ml. After resupplying amino acid-free media with other amino acids (stock solutions^88^), L-cystine stock solution was added to constitute indicated cystine concentration.

### Cystine uptake assay

Cystine uptake was measured by Cystine Uptake Assay Kit (Dojindo). 7×10^5^ cells collected from 1ml culture from a 24-well plate from indicated conditions (RPMI 1640 media with 100 or 1μM methionine for 30 hours, or as a control in 100μM methionine and treated with 1μM erastin for 24 hours) were pelleted and resuspended in 200μl of pre-warmed cystine-free, serum-free media in a U-bottom 96-well plate and incubated at 37°C for 5 minutes in a 5% CO^2^ humidified incubator. Cells were pelleted and resuspended in 200μl of pre-warmed cystine analog uptake solution with fluorescent cystine analog added to recommended concentration and incubated at 37°C for 30 minutes in a 5% CO^2^ humidified incubator. Cells were washed with PBS three times and were extracted for metabolites using 50μl methanol. 200μl working solution prepared accordion to user manual was then added to each well and mixed. After another 30min at 37°C incubation, fluorescence was read with a SpectraMax iD5e plate reader (Molecular Devices). Sample readings were standardized to cells incubated in cystine-free, serum-free medium with no cystine analog but underwent identical treatment.

### NADPH/NADP+ quantitation

NADPH/NADP+ ratios were calculated by measuring NADP and NADPH levels individually. Briefly, cells were resuspended in PBS at ∼1×10^6^/ml and equal amounts were transferred into two wells of a 96-well plate. To measure NADP+ content, cells were lysed in basic solution composed of 200mM NaOH + 1% dodecyltrimethylammonium bromide (Sigma) at 60°C for 15 minutes. To measure NADPH, cells were lysed in acid solution (0.4 N HCl) at 60°C for 15 minutes. Appropriate volumes of Trizma® base (Promega) were added to the basic solution wells and HCl/Trizma® (Promega) the acidic wells according to the manufacturer’s protocol. Luminescence signals were read with a SpectraMax iD5e plate reader. Sample reading were standardized to live cell number determined as described above.

### Mouse xenograft experiments

All animal work was approved by the Cornell University Institutional Animal Care and Use Committee (IACUC) under performed under protocol 2017-0035, current approval 4/22/26.

Dietary methionine restriction in GM12878 cell line derived xenograft (CDX). 32 NSG mice were subcutaneously xenotransplanted in the left flank with finely minced, 2-3 mm^3 pieces of large cell lymphoma GM12878, and placed on a control, methionine-replete diet (TD01084, Envigo) at time of implant. Tumor growth was monitored twice weekly. Once tumor burden reached approximately 30-50 mm^3, as measured by caliper, 24 mice with similar tumor burden were randomized into three different collection timepoint groups; 7, 14 and 21 days (n=8 per timepoint). A tail vein pre-bleed was performed on all 24 mice after which 4 mice from each group were placed on a a 90% reduced methionine diet but that contained balanced total amino acid levels (TD200744, Envigo). Tumor burden and body weight were monitored twice weekly. Tail bleeds continued weekly unless animals were scheduled for collection for which a terminal cardiac bleed was performed. Also at time of sacrifice, tumor was excised and samples were collected either as flash frozen in LN_2_, placed in 10% neutral buffered formalin (moved to 70% EtOH after 24 hours) or placed in RNAlater. All animals were euthanized by CO_2_ asphyxiation at a flow rate 3.5 L/min. All blood was spun for serum and frozen at-20°C and moved to-80°C 24 hours later. Upon inspection of serum metabolomic data, it was evident that serum samples from control and methionine restricted mice had been swapped, and this was corrected.

Methioninease & Cyst(e)inase Administration in GM12878 CDX: 24 NSG mice were subcutaneously xenotransplanted in the left flank with finely minced, 2-3mm^3 pieces of large cell lymphoma GM12878. All mice were fed methionine replete diets. Sixteen mice with similar tumor burden were randomized into four different treatment groups; PBS vehicle, methioninase (90mg/kg), cyst(e)inase (1.5mg/animal), or methioninase (90mg/kg) + cyst(e)inase (1.5mg/animal). Mice were given IP injections on days 16, 18, 20, 22, 24 and 27 post-tumor implantation. Tail bleeds were performed on all study mice before the first treatment and one week later. On Day 28, mice were euthanized by CO_2_ asphyxiation at a flow rate of 3.5L/min at which time a terminal cardiac bleed was performed. Tumor was excised at time of euthanasia and collected three ways; flash frozen in LN2, fixed in 10% neutral buffered formalin (moved to 70% EtOH after 24 hours), and placed in RNAlater. All blood was spun for serum and frozen at - 20°C and moved to-80°C 24hours later. Upon inspection of serum metabolomic data, it was evident that one of the four cyst(e)inase-treated mice had not been successfully dosed, as serum cyst(e)ine levels were similar to PBS-treated mice. Data from this mouse were therefore removed from the analysis.

### Quantification and statistical analysis

For primary B cells, three donors, with three biological replicates from each donor, were used. Statistical details of all experiments can be found in the figure legends. Sample numbers are mentioned in each figure legend and denote biological replicates. Student’s t-test used were 2 tailed and unpaired. Significance among multiple groups was tested using a one-way analysis of variance (ANOVA). For all ANOVAs, the Tukey-Kramer test was used for post hoc multiple comparison, as this allows for the possibility of unequal sample sizes and takes into account the scatter of both groups. GraphPad Prism 11 was used for statistical analysis. Gene ontology analysis was done with the Enrichr module using the KEGG pathway databases. Default Enrichr module parameters were used, with the exception that the Enrichment statistic was set as classic. Metabolic pathway analysis were performed using MetaboAnalyst 5.0.

---

**Figure S1.**
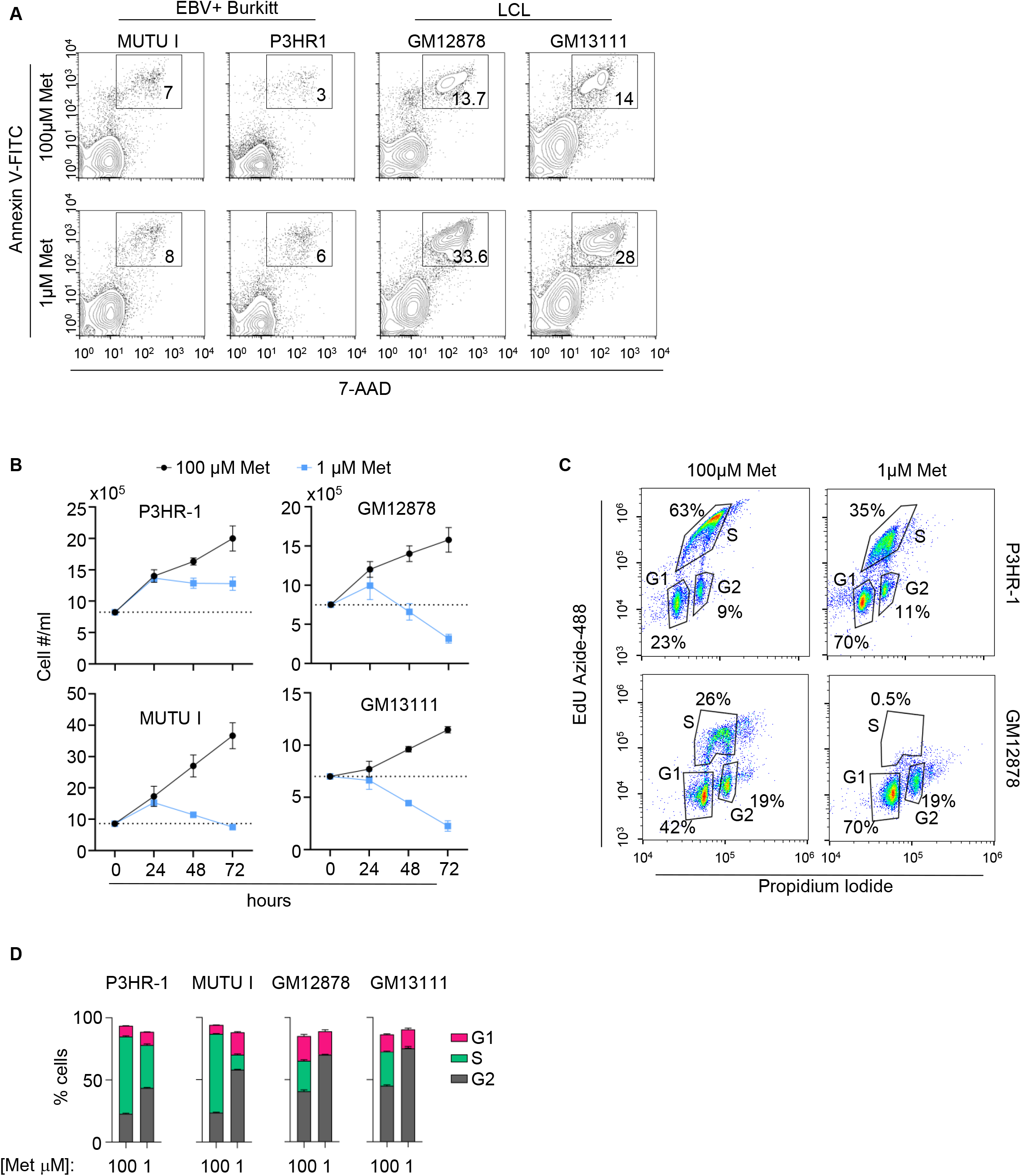
Methionine restriction effects on LCL death or growth arrest, related to. Figure 1. (A) Methionine restriction effects on EBV+ Burkitt versus LCL death. Shown are FACS analysis of EBV+ P3HR-1 or MUTU I Burkitt versus GM12878 or GM13111 LCLs that were cultured in RPMI media containing 100 versus 1μM methionine (met) for 30 hours. Plots are representative of n=3 replicates. (B) Methionine restriction effects on EBV+ Burkitt versus LCL proliferation. Shown are mean ± SD of live cells cultured in media with 100 or 10 μM methionine for the indicated times. (C) Methionine restriction effects on EBV+ Burkitt versus LCL cell cycle. Representative FACS cell cycle analysis from n=3 replicates of P3HR-1 or GM12878 cultured in media with 100 vs 1 μM methionine for 18 hours, using 30 min EdU incorporation and propidium iodide staining. Gates show cells in the G1, S or G2 cell cycle phases. Dead cells were depleted prior to fixation and staining. (D) Methionine restriction effects on EBV+ Burkitt versus LCL cell cycle. Mean ± SD values of G2, S or G1-phase cells from n=3 replicates as in (C). Student’s t-test ****p<0.0001, ***p<0.001, **p<0.01, *p<0.05.

**Figure S2.**
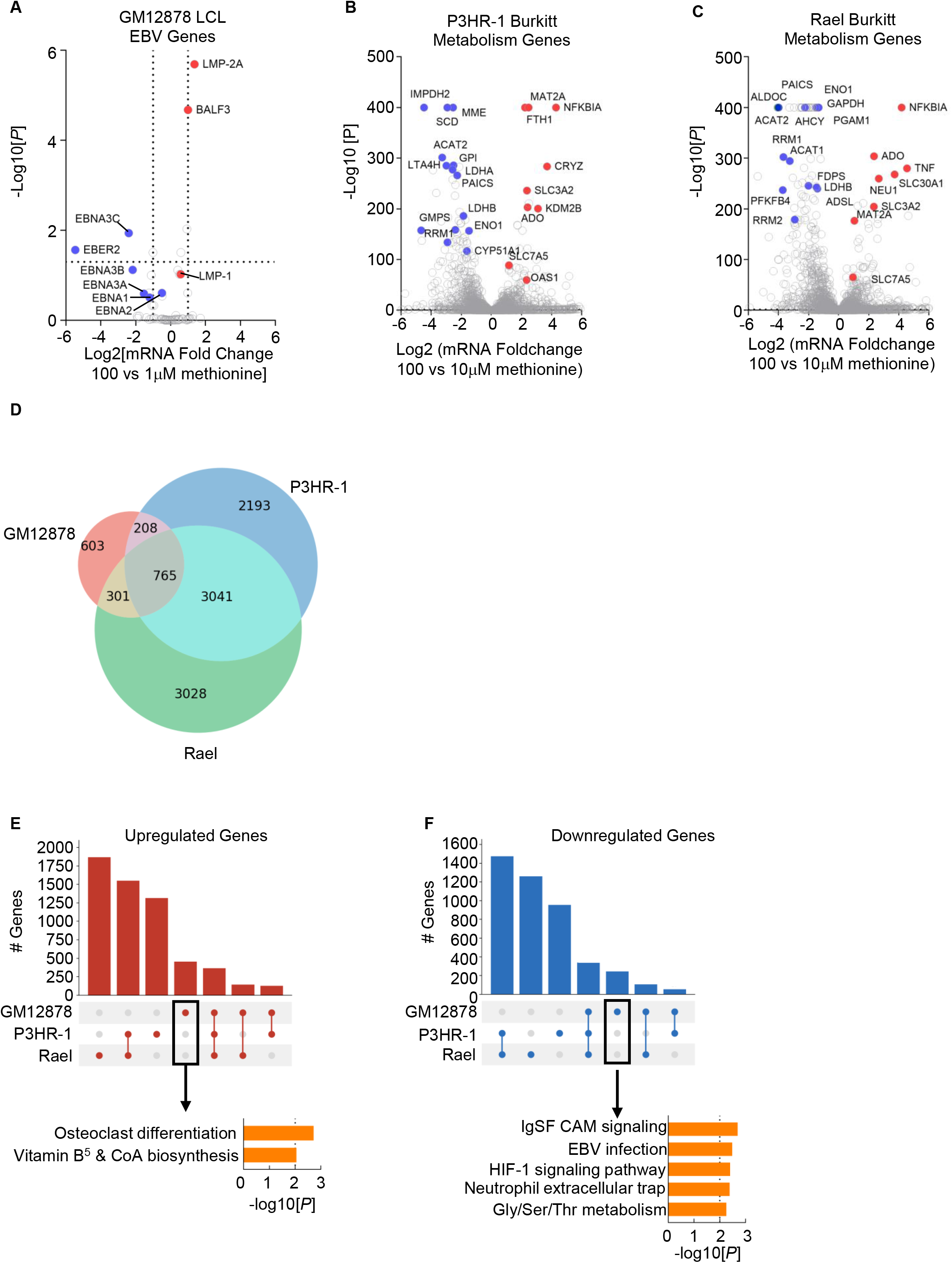
Methionine restriction effects on LCL versus Burkitt gene expression, related to. Figure 1. (A) Methionine restriction effects on GM12878 EBV genes. Volcano plot from n=3 RNAseq replicates of GM12878 grown in media with 100 vs 1μM methionine for 30 hours, as in Fig. 1C. (B) Methionine restriction effects on LCL metabolism gene^89^ expression. Volcano plot analysis of P3HR-1 metabolism gene expression from n=3 RNAseq replicates of P3HR-1 Burkitt cells grown in media with 100 vs 10μM methionine for 72 hours. Data are from^29^. (C) Methionine restriction effects on LCL metabolism gene^89^ expression. Volcano plot analysis of P3HR-1 metabolism gene expression from n=3 RNAseq replicates of EBV+ Rael Burkitt cells grown in media with 100 vs 10μM methionine for 72 hours. Data are from^29^. (D) Venn diagram cross-comparing differentially expressed genes in the indicated cells grown under methionine replete versus restricted conditions (as in A-C), using an adjusted P-value <0.05 cutoff. (E) Cross-comparison of GM12878 LCL versus Burkitt genes upregulated by methionine restriction. Shown are upset plot showing the number of genes commonly upregulated by methionine restriction in the groups indicated below the bar graph by red circles. Shown at bottom are metabolic pathways enriched in genes uniquely upregulated by methionine restriction in GM12878 but not in P3HR-1 or Rael. (F) Cross-comparison of GM12878 LCL versus Burkitt genes downregulated by methionine restriction. Shown are upset plot showing the number of genes commonly downregulated by methionine restriction in the groups indicated below the bar graph by red circles. Shown at bottom are metabolic pathways enriched in genes uniquely downregulated by methionine restriction in GM12878 but not in P3HR-1 or Rael.

**Figure S3.**
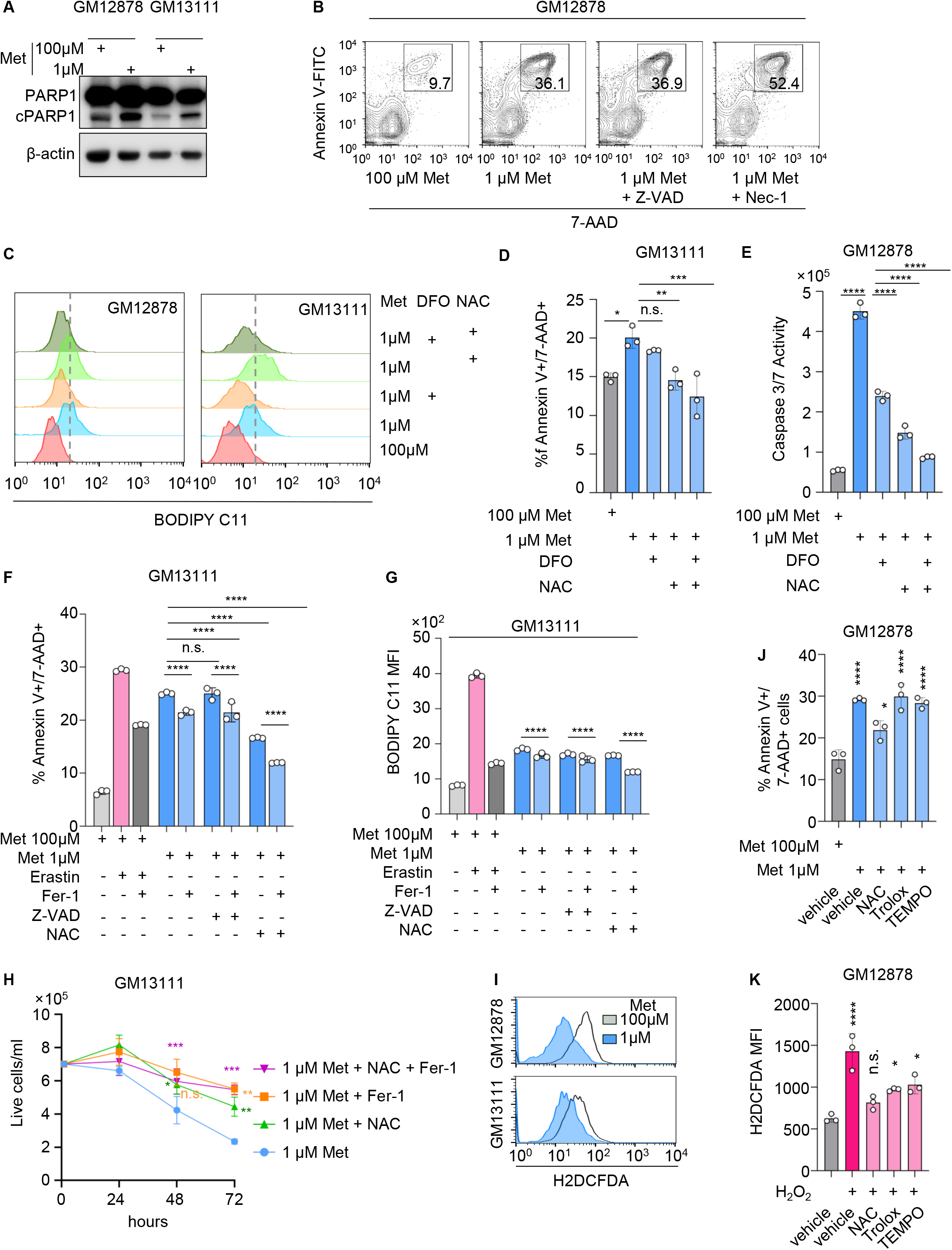
Methionine restriction triggers LCL ferroptosis, related to. Figure 2. (A) Methionine restriction effects on LCL PARP1 cleavage. Immunoblot analysis of whole cell lysates (WCL) from GM12878 or GM13111 LCLs that were cultured in media with 100 vs 1μM methionine for 30 hours. cPARP1, cleaved PARP1. (B) Caspase or RIPK1 inhibition do not rescue methionine restriction induced LCL death. Representative FACS plots from n=3 replicates of annexin-V vs 7-AAD levels in GM12878 LCLs cultured in 100 vs 1μM methionine, in the absence or presence of Z-VAD or Nec-1 (180μM) for 30 hours, as indicated. (C) DFO and NAC effects on LCL BODIPY C11 levels. Representative FACS plots from n=3 replicates of LCLs grown in media with 100 vs 1μM methionine and with DFO and/or NAC for 30 hours, as indicated. (D) DFO and NAC effects on methionine restriction induced GM13111 LCL death. Mean ± SD dead cell (7-AAD+/Annexin V+ cell) percentages from n=3 replicates of GM13111 cultured in media with 100 vs 1μM methionine with DFO and/or NAC for 30 hours, as indicated. (E) Iron chelation and antioxidant effects on methionine-restriction driven caspase activity. Mean ± SD caspase 3/7 values from n=3 replicates of GM12878 LCLs cultured in media with 100 vs 1μM methionine and treated with deferoxamine (DFO, 100μM) and/or N-acetyl cysteine (NAC, 10mM) for 48 hours. (F) Antioxidant versus caspase inhibitor effects on methionine restriction induced GM13111 LCL death. Mean ± SD 7-AAD+/Annexin V+ cell percentages from n=3 replicates of GM13111 cultured in media with 100 vs 1μM methionine and treated with Fer-1, NAC and/or Z-VAD-FMK for 30 hours, as indicated. (G) Antioxidant versus caspase inhibitor effects on methionine restriction induced GM13111 lipid ROS levels. Mean ± SD BODIPY C11 levels from n=3 replicates of GM13111 cultured as in (F). (H) Antioxidant rescue of methionine restriction driven GM13111 LCL death. Mean ± SD live cell numbers from n=3 replicates of GM13111 cultured in media with 1μM methionine and treated with Fer-1 and/or NAC for 30 hours, as indicated. (I) Methionine restriction effects on LCL intracellular oxidant levels. Representative FACS plots from n=3 replicates of GM12878 or GM13111 LCLs cultured in media with 100 or 1μM methionine for 30 hours and stained by H2DCFDA. (J-K) Antioxidant effects on LCL methionine restriction induced LCL death. (J) Mean ± 7-AAD+/Annexin V+ cell percentages from n=3 replicates of GM12878 LCLs cultured in media with 100 vs 1μM methionine and treated with NAC (10 mM), Trolox (50μM) or TEMPO (12.5μM), for 30 hours. (K) As a positive control, shown are mean ± SD H2DCFDA MFI levels from n=3 replicates of GM12878 cultured in media with 100μM methionine and treated with hydrogen peroxide (H_2_0_2_, 50μM), alone or together with NAC (10 mM), Trolox (50μM) or TEMPO (12.5μM), for 30 hours, as indicated and stained by H2DCFDA prior to FACS analysis. Student’s t-test was used for (A). One-way ANOVA was used for (D, E, J, K, and H). Two-way ANOVA was used for F and G. For all statistical analyses, ****p<0.0001, ***p<0.001, **p<0.01, *p<0.05.

**Figure S4.**
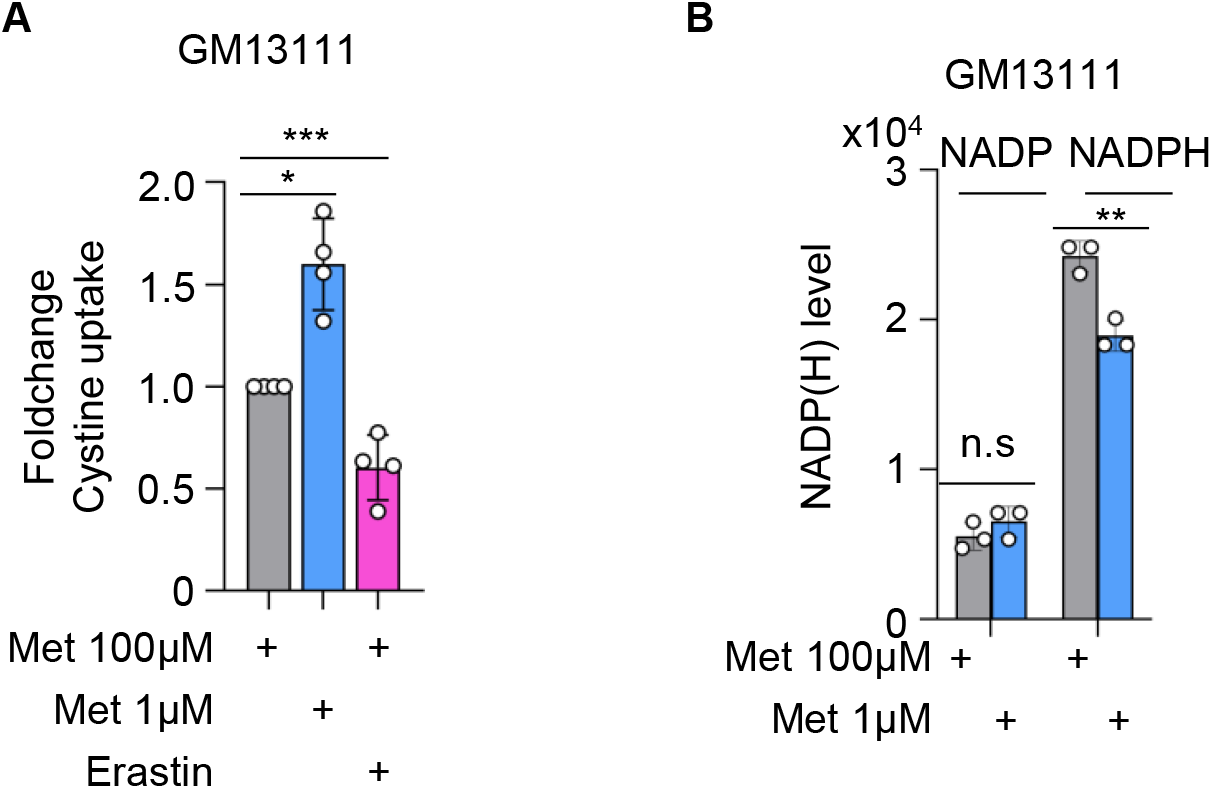
Methionine restriction effects on GM13111 LCL cystine uptake, NADP and NADPH levels, related to. Figure 4. (A) Methionine restriction effects on GM13111 LCL cystine uptake. Mean ± SD foldchange of cystine uptake in levels from n=3 replicates of GM13111 cultured in media with 100 or 1μM Met for 30 hours, or as a control grown in 100μM Met and treated with erastin (1μM) for 18 hours. Cystine uptake was determined by a platereader luminescence assay. (B) Methionine restriction effects on GM13111 LCL NADP(H). Mean ± SD relative NADP(H) levels in GM13111 cultured in media with 100 or 1μM Met, as measured by a NADP(H) luminescence assay. One-way ANOVA was used in (A). Student’s t-test was used in (B). ***p<0.001, **p<0.01, *p<0.05, ns, non-significant.

**Figure S5.**
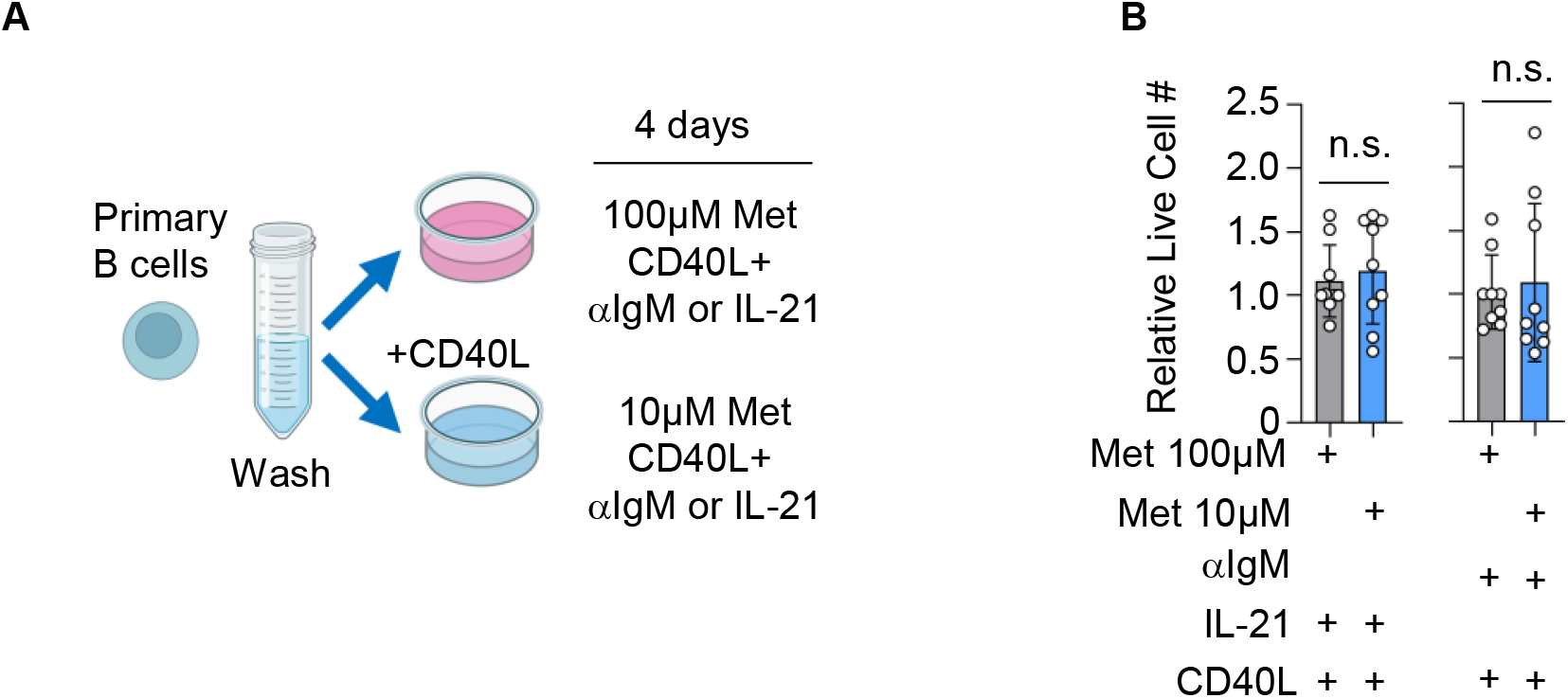
B cell immunoreceptor signaling does not induce peripheral blood B cell methionine survival dependence, related to. Figure 5. (A) Experimental scheme. Peripheral blood B cells cultured in media with 100 vs 10μM Met were stimulated with CD40L (10ng/ml) together with αIgM (1μg/ml) or IL-21 (100ng/ml) for four days. (B) Immunoreceptor signaling effects on methionine survival dependence. Mean ± SD relative live cell numbers of peripheral blood B cells from n=3 donors (3 replicates each) treated as in Student’s t test was used for statistical tests in (B). ns, non-significant.

**Figure S6.**
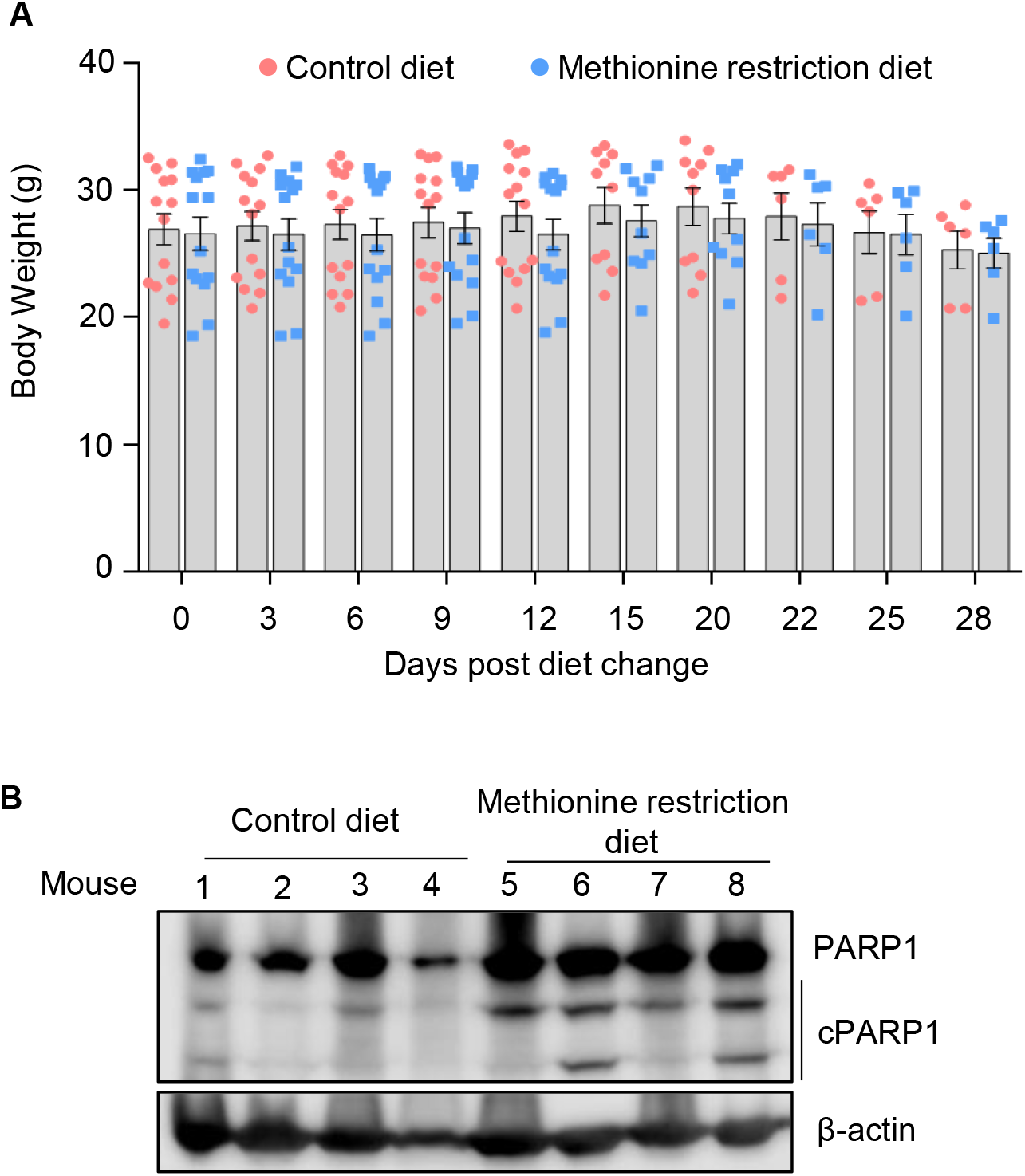
Dietary methionine restriction effects on NOD-SCID mice and LCL xenografts, related to. Figure 6. (A) Dietary methionine restriction effects on murine body weight. Mean ± SD body weight measurements for mice fed control (red) or methionine restriction (blue) diets. (B) Dietary methionine restriction effects on cleaved PARP levels. Representative immunoblot from n=3 replicates of WCL from LCL xenografts explanted at day14.

**Figure S7.**
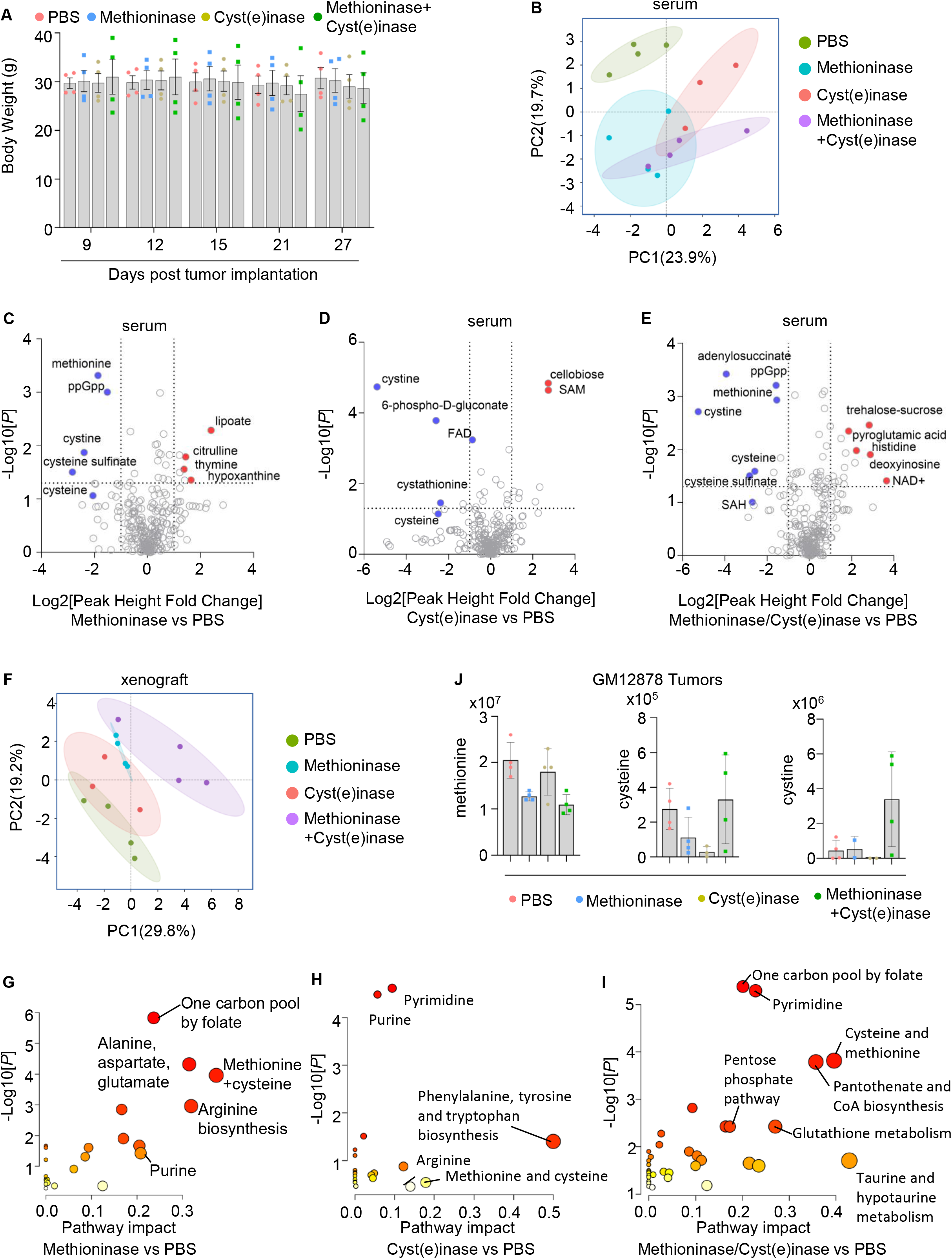
Methioninase and cyst(e)inase effects on GM12878 xenografts in vivo, related to. Figure 7. (A) Methioninase and cyst(e)inase effects on mouse body weight. Mean ±SD body weight measurements at the indicated days post-xenograft implantation of mice administered PBS, methioninase, cyst(e)inase or both from the mice treated in Fig. 7. (B) PCA plot of serum amino acid metabolites (KEGG ID:ko00270) analysis from the mice treated in Fig. 7. Serum was collected at day 28 post-tumor implantation. (C) Methioninase effects on NOD-SCID serum metabolomes. Volcano plot from LC-MS polar metabolite analysis of n=4 LC-MS replicates of serum samples collected at day 28 from mice that were administered PBS versus methioninase. (D) Cyst(e)inase effects on NOD-SCID serum metabolomes. Volcano plot from LC-MS polar metabolite analysis of n=4 LC-MS replicates of serum samples collected at day 28 from mice that were administered PBS versus cyst(e)inase. (E) Combined methioninase/cyst(e)inase effects on NOD-SCID serum metabolomes. Volcano plot from LC-MS polar metabolite analysis of n=4 LC-MS replicates of serum samples collected at day 28 from mice that were administered PBS versus methioninase + cyst(e)inase. (F) PCA plot of GM12878 tumor amino acid metabolites (KEGG ID:ko00270) analysis from the mice treated in Fig. 7. (G) MetaboAnalyst analysis of methioninase driven metabolic pathway impact on GM12878 xenograft tumors, related to Fig 7C. Higher pathway impact values indicates stronger methioninase effects restriction on the indicated pathway. (H) MetaboAnalyst analysis of cyst(e)inase driven metabolic pathway impact on GM12878 xenograft tumors, related to Fig 7D. Higher pathway impact values indicates stronger methioninase effects restriction on the indicated pathway. (I) MetaboAnalyst analysis of methioninase/cyst(e)inase driven metabolic pathway impact on GM12878 xenograft tumors, related to Fig 7E. Higher pathway impact values indicates stronger methioninase effects restriction on the indicated pathway. (J) Methioninase and/or cyst(e)inase effects on LCL tumor methionine, cysteine or cystine levels. Mean ± SD tumor methionine, cysteine or cystine ion intensities from Figure 7 LC-MS analyses. Student’s t-test was used for statistical analyses, **p<0.01, *p<0.05. ns, non-significant.

