## Supplementary material for "Epstein-Barr virus transformation creates a methionine-dependent ferroptosis vulnerability in B cells": Reagents and Resources

**Key resources table**

| REAGENT or RESOURCE | SOURCE | IDENTIFIER |
| --- | --- | --- |
| Antibodies | | |
| Anti-β-actin Antibody | Biolegend | 664802; RRID: AB_2721349 |
| Anti-phoshpo-MLKL (Thr357/Ser358) rabbit antibody | Cell signaling | 14516 |
| PARP rabbit monoclonal antibody (clone 46D11) | Cell Signaling Technology | 9532S |
| Phospho-MLKL (Thr357/Ser358) rabbit monoclonal antibody (clone D6H3V) | Cell Signaling Technology | 14516S |
| CBS (D8F2P) Rabbit mAb #14782 | Cell Signaling | 14782s |
| Cystathionine γ-Lyase (D4E9J) Rabbit mAb #30068 | Cell Signaling | 30068S |
| Mouse monoclonal anti-EBV LMP1 antibody (S12) | A gift from Dr. David Thorley-Lawson | N/A |
| EBV LMP2a rat monoclonal antibody | Santa Cruz | sc-101314 |
| Mouse monoclonal anti-EBNA2 antibody (clone PE2) | A gift from Dr. Elliott Kieff | N/A |
| Mouse monoclonal anti-EBNA3A antibody (clone A10) | A gift from Dr. Elliott Kieff | N/A |
| Sheep anti EBNA3A, pAb | Exalpha | F115P |
| Phospho-ATM (Ser1981) rabbit monoclonal antibody (clone D25E5) | Cell Signaling Technology | 13050 |
| Hyperoxidized peroxiredoxin-3 monoclonal antibody (clone 5H7c) | Cayman Chemical | 39888 |
| Anti-rabbit IgG, HRP-linked Antibody | Cell Signaling Technology | 7074V |
| Anti-mouse IgG, HRP-linked Antibody | Cell Signaling Technology | 7076V |
| Anti-rat IgG, HRP-linked Antibody | Cell Signaling Technology | 7077S |
| Anti-Goat IgG, HRP-linked Antibody | Thermo Fisher | #A24452 |
| Bacterial and virus strains | | |
| B95.8 EBV | A gift from Dr. Elliott Kieff | N/A |
| B95.8 WT BAC | A gift from Dr. Wolfgang Hammerschmidt^1^ | N/A |
| B95.8 ΔEBNA2 BAC | A gift from Dr. Wolfgang Hammerschmidt^1^ | N/A |
| B95.8 ΔLMP1 BAC | A gift from Dr. Wolfgang Hammerschmidt^1^ | N/A |
| B95.8 ΔLMP2A BAC | A gift from Dr. Wolfgang Hammerschmidt^1^ | N/A |
| B95.8 ΔEBNALP BAC | A gift from Dr. Wolfgang Hammerschmidt^1^ | N/A |
| B95.8 ΔEBNA3A BAC | A gift from Dr. Wolfgang Hammerschmidt^1^ | N/A |
| B95.8 ΔEBNA3C BAC | A gift from Dr. Wolfgang Hammerschmidt^1^ | N/A |
| Chemicals, peptides, and recombinant proteins | | |
| L-Lysine monohydrochloride | Thermo Fisher | A16249.18 |
| L-Arginine | Thermo Fisher | A15738.22 |
| L-Histidine | Millipore Sigma | H8000 |
| L-Phenylalanine | Thermo Fisher | J63925.22 |
| L-Serine | Thermo Fisher | J62187.09 |
| L-Threonine | Millipore Sigma | T8625 |
| L-Tryptophan | Thermo Fisher | J62508.09 |
| L-Tyrosine disodium salt dihydrate | Thermo Fisher | J61770.22 |
| L-Hydroxyproline | ChemScene | CS-W008928 |
| L-Leucine | Millipore Sigma | L8000 |
| L-Isoleucine | Millipore Sigma | I2752 |
| L-Valine | Millipore Sigma | 94619 |
| L-Glutamic acid | Millipore Sigma | G1251 |
| L-Methionine | Millipore Sigma | M5308 |
| L-Asparagine | Millipore Sigma | A4159 |
| L-Aspartic acid | Millipore Sigma | A9256 |
| L-Cystine | Millipore Sigma | C8755 |
| L-Proline | Millipore Sigma | P5607 |
| Glysine | Thermo Fisher | AC120070050 |
| Gibco™ L-Glutamine (200 mM) | Life Technologies | 25030-164 |
| Trypan Blue Solution, 0.4% | Thermo Fisher | 15250061 |
| Water (LC/MS) | Fisher Scientific | Cat#W6-4 |
| Formaldehyde solution | Sigma-Aldrich | F8775 |
| ProLong™ Gold Antifade Mountant with DAPI | Thermo Fisher | P36935 |
| Formaldehyde solution | Sigma-Aldrich | F8775 |
| MegaCD40L, Soluble (human) (recombinant) | Enzo life sciences | ALX-522-110-C010 |
| Recombinant Human IL-21 (carrier-free) | Biolegend | 571208 |
| Anti-Human IgM (μ-chain specific) antibody produced in goat | Sigma | I0759 |
| 5-Ethynyl-2'-deoxyuridine (EdU) | MCE | HY-118411 |
| RNase A, DNase and protease-free (10 mg/mL) | Thermo Fisher | EN0531 |
| TransIT-LT1 Transfection Reagent | Mirus | MIR 2306 |
| BSA, Molecular Biology Grade | NEB | B9000S |
| ECL chemiluminescence | Millipore Sigma | WBLUF0500 |
| Z-VAD-FMK | Selleck | S7023 |
| Necrostatin-1 - RIPK1 Inhibitor | InvivoGen | inh-ncst1 |
| L-methionine (13C5, 99%) | Cambridge Isotope Lab | CLM-893-H-PK |
| β-cyano-L-alanine | Cayman | 10010947 |
| DL-Homocysteine thiolactone hydrochloride | Thermo Fisher Scientific | L09077.22 |
| DL-Propargyl Glycine (hydrochloride) | Cayman | 10010948 |
| cysteamine | Sigma | M9768 |
| Puromycin Dihydrochloride | Thermo Fisher Scientific | A1113803 |
| PEI MAX | Polysciences | 24765-1 |
| Critical commercial assays | | |
| Invitrogen PureLink Quick Plasmid Maxiprep Kit | Invitrogen | K210016 |
| Pierce™ BCA Protein Assay Kit | Thermo Fisher | 23225 |
| QIAprep Spin Miniprep Kit | Qiagen | 27106 |
| NEBNext® Poly(A) mRNA Magnetic Isolation Module | New England Biolabs | E7490S |
| NEBNext® Ultra™ II Directional RNA Library Prep with Sample Purification Beads | New England Biolabs | E7765S |
| NEBNext® Multiplex Oligos for Illumina® (Index Primers Set 2) | New England Biolabs | E7500S |
| NEBNext® Multiplex Oligos for Illumina® (Index Primers Set 1) | New England Biolabs | E7335S |
| Caspase-Glo® 3/7 Assay | Promega | G8201 |
| BODIPY™ 581/591 C11 | Invitrogen | D3861 |
| Cystine Uptake Assay Kit | Dojindo | up05-12 |
| NADP/NADPH-glo assay | Promega | G9071 |
| H2DCFDA | Life Technologies | D399 |
| FITC-conjugated Annexin V antibody | Biolegend | 640945 |
| 7-AAD (7-Aminoactinomycin D) | ThermoFisher Scientific | Cat#A1310 |
| Alexa Fluor™ 488 Azide | Thermo Fisher | A10266 |
| Propidium Iodide - 1.0 mg/mL Solution in Water | Thermo Fisher | P3566 |
| Dodecyltrimethylammonium bromide (DTAB) | Sigma | D8638-25G |
| Deposited data | | |
| GM12878 RNA seq | This paper | GEO GSE336950 |
| Experimental models: Cell lines | | |
| EBV+ Burkitt lymphoblastoid cell GM12878-Cas9 | Ma et al., 2017^2^ | N/A |
| EBV+ Burkitt lymphoblastoid cell GM13111-Cas9 | Ma et al., 2017^2^ | N/A |
| EBV+ Burkitt lymphoma P3HR1-Cas9 | Ma et al., 2017^2^ | N/A |
| 293T | ATCC | CRL-3216 |
| EBV+ Burkitt lymphoma MUTU I | Dr. Jeffrey Sample | N/A |
| EBV+ Burkitt lymphoma MUTU III | Dr. Jeffrey Sample | N/A |
| EBV- Burkitt lymphoma Raji | Dr. Elliot Kieff | N/A |
| Experimental models: Organisms/strains | | |
| NSG (NOD/SCID gamma) humanized mice | The Jackson Laboratory | 5557 |
| Recombinant DNA | | |
| 509 MacVec | A gift from Dr. Wolfgang Hammerschmidt | N/A |
| 2670_(pRA) | A gift from Dr. Wolfgang Hammerschmidt | N/A |
| Software and algorithms | | |
| Image Studio Ver 5.5 | Li-COR | https://www.licor.com/bio/image-studio/ |
| GraphPad Prism 11 | GraphPad Software | https://www.graphpad.com/scientific-software/prism/ |
| FlowJo V10 | Flowjo LLC. | https://www.flowjo.com/ |
| MetaboAnalyst 6.0 | Wishart Research Group | www.Metaboanalyst.ca |
| FIJI ImageJ | Open Source | imagej.net/software/fiji |
| STAR v2.7.10a | Open Source | https://github.com/alexdobin/STAR |
| DESeq2 v1.48.2 | Open Source | https://bioconductor.org/packages/release/bioc/html/DESeq2.html |
| Biorender | Biorender | https://biorender.com/ |
| ggplot2 | Open Source | https://ggplot2.tidyverse.org |
| Other | | |
| Standard Fetal Bovine Serum, Qualified, USDA-Approved Regions | ThermoFisher Scientific | 10437028 |
| Dialyzed Fetal Bovine Serum | ThermoFisher Scientific | 26400044 |
| DMEM, high glucose, pyruvate | Life Technologies | 11875135 |
| RPMI 1640 Medium | Life Technologies | 11995081 |
| RPMI 1640 medium without amino acids, sodium phosphate | Thermo Fisher | R8999-04A |
| RPMI, no methionine | ThermoFisher Scientific | A1451701 |
| RosetteSep Human B Cell Enrichment Cocktail | STEMCELL Technologies | Cat#15064 |
| EasySep Human B Cell Enrichment Kit | STEMCELL Technologies | Cat#19054 |
| High Precision Glass Cover Slip, box of 100, No 1.5, 24x50mm | Bioscience Tools | Cat#CSHP-No1.5-24x50 |
| Methionine replete diet (Control diet) | Envigo | TD.01084 |
| Methionine restricted diet (0.084% Met diet) | Envigo | TD.200744 |
| Opti-MEM™ I Reduced Serum Medium | ThermoFisher | 31985062 |

1.	Pich, D., Mrozek-Gorska, P., Bouvet, M., Sugimoto, A., Akidil, E., Grundhoff, A., Hamperl, S., Ling, P.D., and Hammerschmidt, W. (2019). First Days in the Life of Naive Human B Lymphocytes Infected with Epstein-Barr Virus. mBio *10*. 10.1128/mBio.01723-19.

2.	Ma, Y., Walsh, M.J., Bernhardt, K., Ashbaugh, C.W., Trudeau, S.J., Ashbaugh, I.Y., Jiang, S., Jiang, C., Zhao, B., Root, D.E., et al. (2017). CRISPR/Cas9 Screens Reveal Epstein-Barr Virus-Transformed B Cell Host Dependency Factors. Cell Host Microbe *21*, 580–591 e587. 10.1016/j.chom.2017.04.005.
